# SARS-CoV-2 evolution in animals suggests mechanisms for rapid variant selection

**DOI:** 10.1101/2021.03.05.434135

**Authors:** Laura Bashor, Roderick B. Gagne, Angela Bosco-Lauth, Richard Bowen, Mark Stenglein, Sue VandeWoude

## Abstract

SARS-CoV-2 spillback from humans into domestic and wild animals has been well-documented. We compared variants of cell culture-expanded SARS-CoV-2 inoculum and virus recovered from four species following experimental exposure. Five nonsynonymous changes in nsp12, S, N and M genes were near fixation in the inoculum, but reverted to wild-type sequences in RNA recovered from dogs, cats and hamsters within 1-3 days post-exposure. Fourteen emergent variants were detected in viruses recovered from animals, including substitutions at spike positions H69, N501, and D614, which also vary in human lineages of concern. The rapidity of in vitro and in vivo SARS-CoV-2 selection reveals residues with functional significance during host-switching, illustrating the potential for spillback reservoir hosts to accelerate evolution, and demonstrating plasticity of viral adaptation in animal models.

**One-Sentence Summary:** SARS-CoV-2 variants rapidly arise in non-human hosts, revealing viral evolution and potential risk for human reinfection.

---

Cross-species transmission events, which challenge pathogens to survive in new host environments, typically result in species-specific adaptations (*1*). These evolutionary changes can determine the pathogenicity and transmissibility of the virus in novel host species (*2*). Cross-species transmission events are rare, and in natural settings typically fail to result in epidemic spread (*3*). In contrast to most species, humans move globally and regularly come into contact with domestic and peridomestic animals. Thus, when a novel virus spreads through human populations, there is an incidental risk of exposure to potentially susceptible non-human species. This scenario has become evident with the SARS-CoV-2 pandemic. Originally resulting from viral spillover into humans (*4,5*), likely from an animal reservoir, spillback into a wide range of companion and wild animals has occurred or been shown to be plausible (*6*-*9*).

Repeated interspecies transmission of a virus presents the potential for acceleration of viral evolution and a possible source of novel strain emergence. This was demonstrated by reverse zoonosis of SARS-CoV-2 from humans to mink, followed by selection in mink and zoonotic transmission back to humans (*8*). Given that reverse zoonosis has been reported repeatedly in dogs and cats from households where COVID-19 patients reside, and the fact that up to 50% of households worldwide are inhabited by these companion animals, there is potential for similar transmission chains to arise via humans and their pets (*10,11*). Elucidating viral selection and species-specific adaptation of SARS-CoV-2 in common companion animals is therefore of high interest. Furthermore, understanding viral evolutionary patterns in both companion animals and experimental animal models provides a valuable appraisal of species-specific viral variants that spotlight genomic regions for host-virus interaction.

Significant attention has been directed at substrains evolving from the initial SARS-CoV-2 isolate (*12*) and an accumulating number of variant lineages have demonstrated increased transmission potential in humans (*13,14*). The role, if any, that reverse zoonotic infections of nonhuman species and spillback may have played in the emergence of these novel variants of SARS-CoV-2 remains unknown. Documenting viral evolution following spillover of SARS-COV-2 into new species is difficult given the unpredictability of timing of these events; therefore, experimental studies can greatly aid understanding of SARS-CoV-2 evolution in animal species. Laboratory-based studies also provide the opportunity to determine how changes that occur during viral expansion in cell culture may influence in vivo infections. This information is highly relevant for interpretation of in vivo and in vitro experiments using inoculum propagated in culture.

We therefore set out to assess the evolution of SARS-CoV-2 during three rounds of expansion of strain USA-WA1/2020 in Vero E6 cells (*15*), followed by changes occurring during primary experimental infection in four mammalian hosts. Specifically, we compared variant proportions, insertions, and deletions occurring in genomes of SARS-CoV-2 recovered from dogs (n=3), cats (n=6), hamsters (n=3) and ferret (n=1).

## Results

### SARS-CoV-2 genome sequence recovery

We recovered full genome sequences of SARS-CoV-2 from stocks of three serial passages of USA-WA1/2020 inoculum and virus recovered from five cats with primary exposure, one cat exposed by contact to an infected cat (*6*), three dogs (*6*), three hamsters, and one ferret (table S1). We obtained a median depth of coverage of 2859x (mean = 4380x) using ARTIC version 3 primers as described in methods (fig. S1). SARS-CoV-2 genomes recovered from Cat 5 and one technical replicate of Cat 4 were amplified with ARTIC version 2 primers with median depth of coverages of 506x and 1056x respectively.

### Identification of single nucleotide and structural variants

Eight-eight unique single nucleotide (SNV) and structural variant (SV; insertions and deletions) were present in 3-100% of viral genome sequences in two technical replicates, and were observed 270 times across all datasets (242 SNV, 28 SV; **Fig. 1A**). This included 74 SNVs and 14 SVs. Seventy-one of 270 variants (26.3%) were detected at 25% frequency or greater (**Fig. 1B**). Seven SARS-CoV-2 SNV recovered from one or more species occurred at positions that coincide with variants of concern in humans or other species (**Table 1**).

**Table 1.**
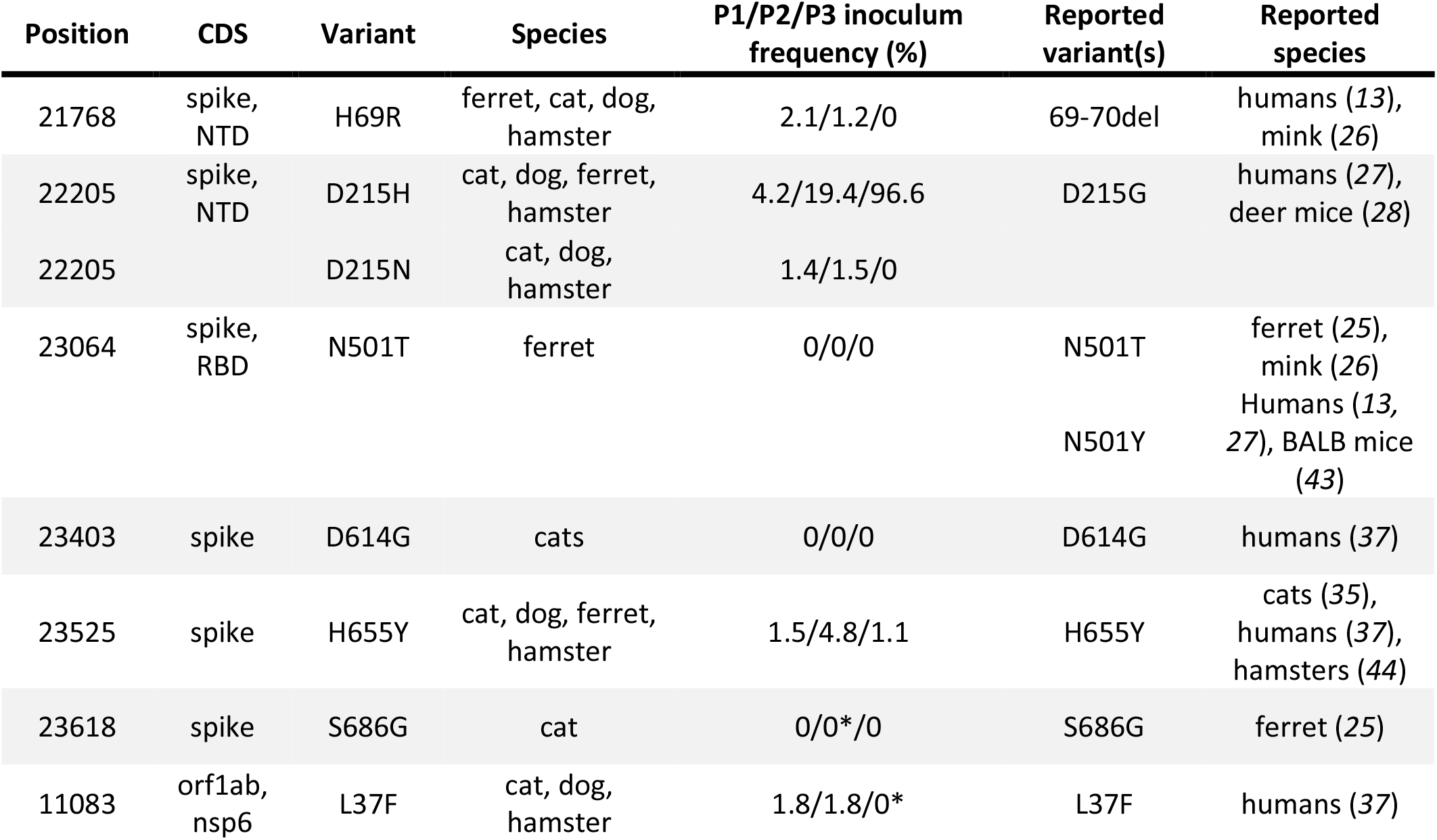
Seven SARS-CoV-2 SNV recovered from one or more species have been reported as variants of concern in humans or other species. Frequency of each variant detected in Vero E6 cell supernatant at each passage (P1, P2, P3) represents the average of two replicates; values undetected above 0.1% are reported as 0. Only D215H was detected at high frequency in P3 inoculum. *S686G was detected in one replicate of P2 at 0.8% and undetected in the second; L37F was detected in one replicate of P3 at 0.5% and undetected in the second.

**Fig 1.**
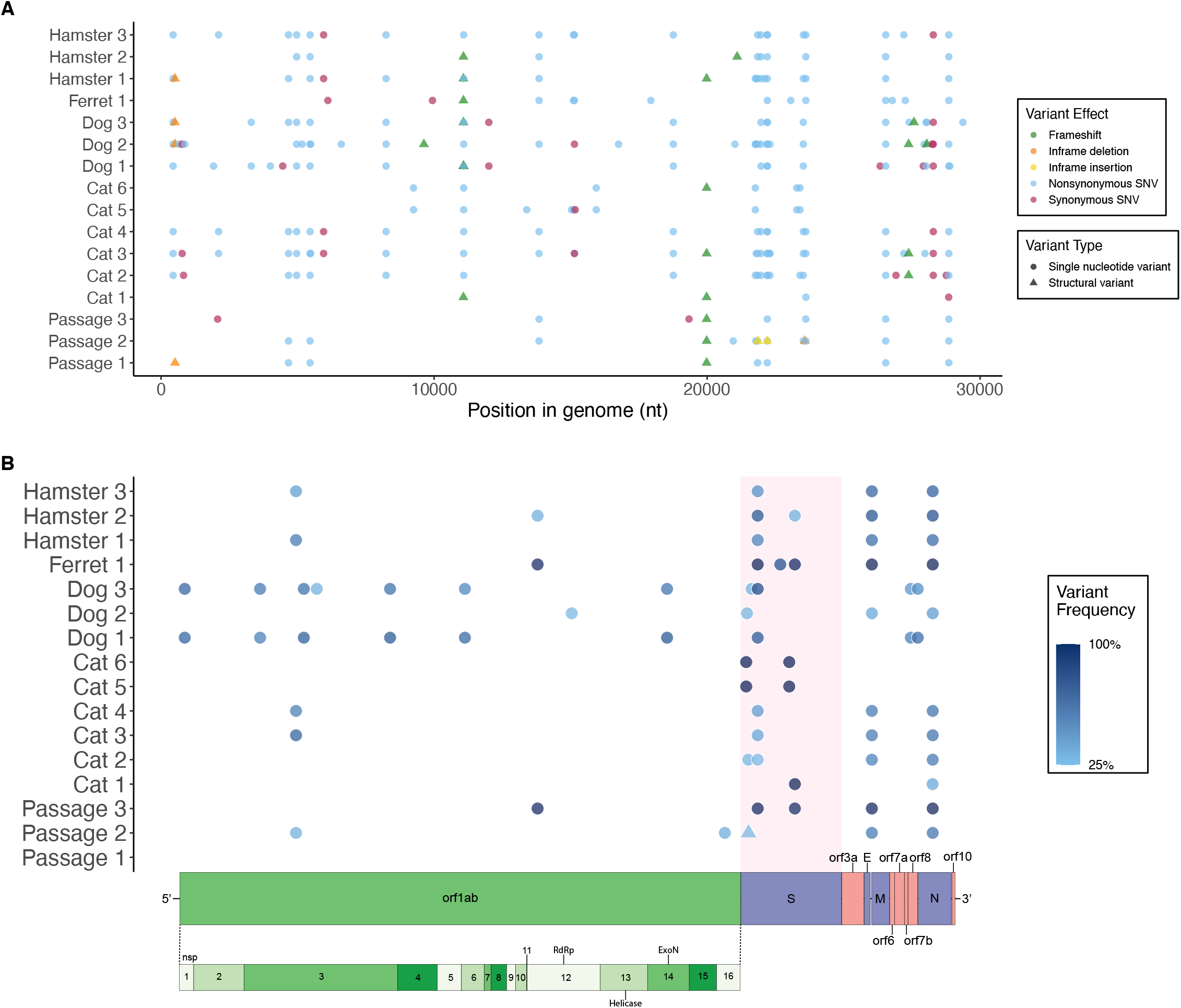
Eight-eight unique variants were detected in >3% of sequences of in vitro-and in vivo-derived SARS-CoV-2. (**A**) Position, predicted effect and (**B**) allele frequency of single nucleotide (SNV) and structural (SV) variants detected across the SARS-CoV-2 genome in sequences obtained from 13 experimentally inoculated animals and three passages of the viral inoculum. Each point represents a SNV (circle) or SV (triangle). (A) All variants detected in ≥3% of sequences, demonstrating a majority of SNVs and a slightly increased occurrence of modifications in the spike protein. (B) All variants detected in ≥ 25% of sequences, revealing the presence of higher frequency variants in the spike protein of all datasets excluding the Passage 1 stock virus, and the prevalence of higher frequency variants across the entire genome of SARS-CoV-2 recovered from dogs. Frequency indicated by darkness scale. Schematic of SARS-CoV-2 genome illustrated for orientation.

### Cell culture-associated viral variants and reversion in animals

Passage of SARS-CoV-2 in Vero E6 cells resulted in five nonsynonymous amino acid changes that reached fixation or near fixation following three passages (**Fig. 2**). These variants reverted to wild type sequences within 1-3 days after inoculation in dogs, cats and hamsters. Reversion frequency was higher in samples recovered three days post-infection (dogs and cats) compared to virus recovered one day post-infection (hamsters, **Fig. 2**). Spike variant D215H underwent a second substitution to D215N, which reached near fixation (>72%) in two dogs. This variant was also detected at >6% frequency in the third dog, one hamster and three cats.

**Fig 2.**
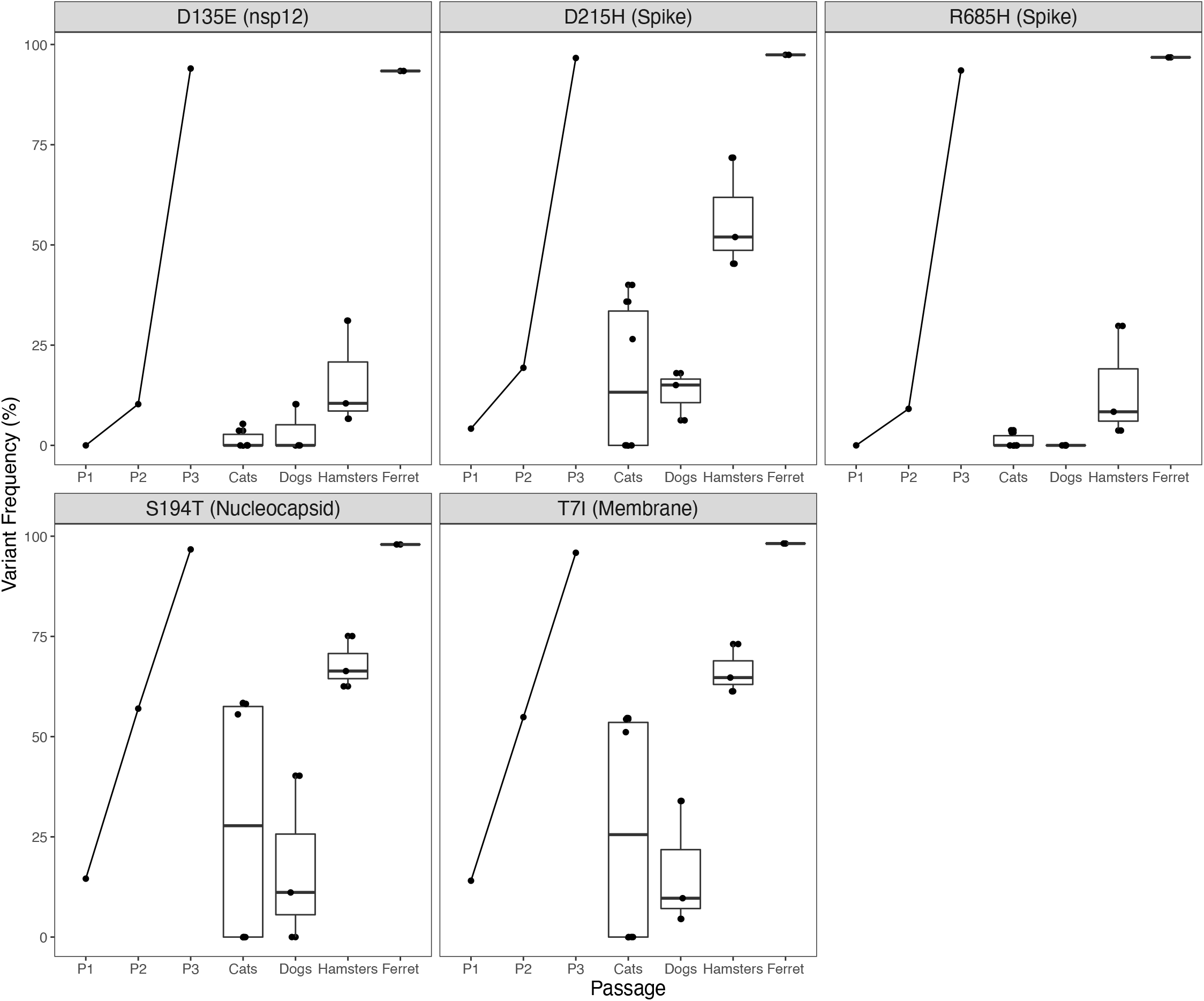
SARS-CoV-2 cell culture variants revert rapidly during in vivo experimental infection. SARS-CoV-2 isolate USA-WA1/2020 was passaged three times in Vero E6 cell line. Five single nucleotide variant substitutions across the genome reached >93% frequency; variant proportion recovered from each supernatant stock indicated by P1, P2, P3. Cats, dogs, hamsters and ferret (n=6, 3, 3, and 1, respectively) were inoculated with 105 – 106 pfu intranasally. Virus was recovered 1-3 days PI, sequenced using a tiled amplicon technique and analyzed with a pipeline for calling single nucleotide and structural variants in viral populations. Cell culture variants decreased in frequency in all animals with the exception of the ferret (n=1). Variants are indicated in reference to the consensus residue in USA-WA1/2020 and their position within the coding sequence of a SARS-CoV-2 protein. Each point represents the mean of two technical replicates, aside from one cat for which a replicate was not sequenced.

### Viral variants arising during in vivo passage

Thirteen nonsynonymous variants and one synonymous variant not present in the USA-WA1/2020-derived viral inoculum were detected at >50% frequency in one individual, or were detected in all individuals of one species (**Table 2**). The default limit of detection for variant calling was set at 3% but it was possible to observe variants with frequencies down to 0.1% with possibly reduced quantitative accuracy (*16*). Variants detected at lower frequencies within the Passage 3 (P3) inoculum in matched replicates would be suggestive of in vivo selection of these pre-existing quasispecies. Using this reasoning, we detected 10 of the 14 emergent variants from animal hosts at 0.1-3% in the viral inoculum, and four were not identified in the viral inoculum at all.

**Table 2.** Nonsynonymous SARS-CoV-2 variants emerge following experimental inoculation in cats, dogs, hamsters and ferret. SARS-CoV-2 isolate USA-WA1/2020 was expanded for three passages in Vero E6 cells and 10^5^–10^6^ pfu was inoculated intranasally in cats, dogs, hamsters and ferret (n=6, 3, 3, and 1). Virus was recovered 1-3 days PI (dog and cat infections described in (*6*)), sequenced using a tiled amplicon technique and analyzed with a pipeline for calling single nucleotide and structural variants in viral populations. Variants representing >50% of recovered genomes in at least one individual or that were found in all individuals of a species are indicated here. Shading indicates species: Cats=yellow; dogs=green; hamster=blue; ferret unshaded. Bold indicates variants occurring in >50% of genomes; boxes indicate variants detected in all sampled individuals of that species. Only one synonymous mutation met the criteria and is displayed. Variants that emerged following passage in Vero E6 cells (Fig. 2) are not listed. Each point represents the mean of two technical replicates, aside from Cat 5 for which a replicate was not sequenced.

| Variant | Position | CDS | N or S | Cat<br>1 | Cat<br>2 | Cat<br>3 | Cat<br>4 | Cat<br>5 | Cat<br>6 | Dog<br>1 | Dog<br>2 | Dog<br>3 | Ferret<br>1 | Hamster<br>1 | Hamster<br>2 | Hamster<br>3 |
| --- | --- | --- | --- | --- | --- | --- | --- | --- | --- | --- | --- | --- | --- | --- | --- | --- |
| G59D | 441 | nsp1 | N |  | 0.15 | 0.06 | 0.10 |  |  | 0.66 | 0.14 | 0.66 |  | 0.04 |  | 0.07 |
| E195G | 3303 | nsp3 | N |  |  |  |  |  |  | 0.53 |  | 0.56 |  |  |  |  |
| T749I | 4965 | nsp3 | N |  | 0.16 | 0.06 | 0.10 |  |  | 0.68 | 0.13 | 0.59 |  |  | 0.04 | 0.07 |
| H1841Y | 8240 | nsp3 | N |  | 0.17 | 0.06 | 0.10 |  |  | 0.70 | 0.13 | 0.59 |  | 0.04 |  | 0.07 |
| L37F | 11083 | nsp6 | N |  | 0.15 | 0.06 | 0.09 | 0.05 | 0.06 | 0.64 | 0.10 | 0.49 |  | 0.04 |  | 0.06 |
| I242V | 18763 | nsp14 | N |  | 0.18 | 0.06 | 0.10 |  |  | 0.71 | 0.15 | 0.58 |  |  |  | 0.06 |
| H69R | 21768 | S | N |  | 0.06 |  |  | 1.00 | 0.99 |  | 0.14 |  |  | 0.06 |  |  |
| D138Y | 21974 | S | N |  | 0.07 | 0.06 | 0.05 |  |  | 0.06 | 0.07 | 0.29 |  | 0.10 | 0.12 | 0.10 |
| D215N | 22205 | S | N |  | 0.17 | 0.07 | 0.10 |  |  | 0.72 | 0.14 | 0.78 |  |  |  | 0.06 |
| N501T | 23064 | S | N |  |  |  |  |  |  |  |  |  | 0.76 |  |  |  |
| D614G | 23403 | S | N |  | 0.16 |  |  | 0.99 | 0.98 |  |  |  |  |  |  |  |
| S686G | 23618 | S | N | 0.99 |  |  |  |  |  |  |  |  |  |  |  |  |
| S43Y | 28021 | orf8 | N |  |  |  |  |  |  | 0.53 |  | 0.44 |  |  |  |  |
| N4N | 28285 | N | S |  | 0.15 | 0.06 | 0.11 |  |  | 0.62 | 0.10 | 0.52 |  |  |  | 0.05 |

There was no significant difference between the mean number of unique variants detected in viral genomes recovered from the four species (**Fig. 3A**). SARS-CoV-2 sequences also did not cluster by species (fig. S2). Excluding variants that were detected in the original inoculum sequences at ≥3% frequency, 69.6% of novel variants were found in a single species, 11.6% were found in two species, and 17.4% were found in three species. Only one variant that was not detectable in the inoculum at ≥3% frequency was detected in all four animal species, a single nucleotide insertion at L37 in nsp6 causing a frameshift.

**Fig 3.**
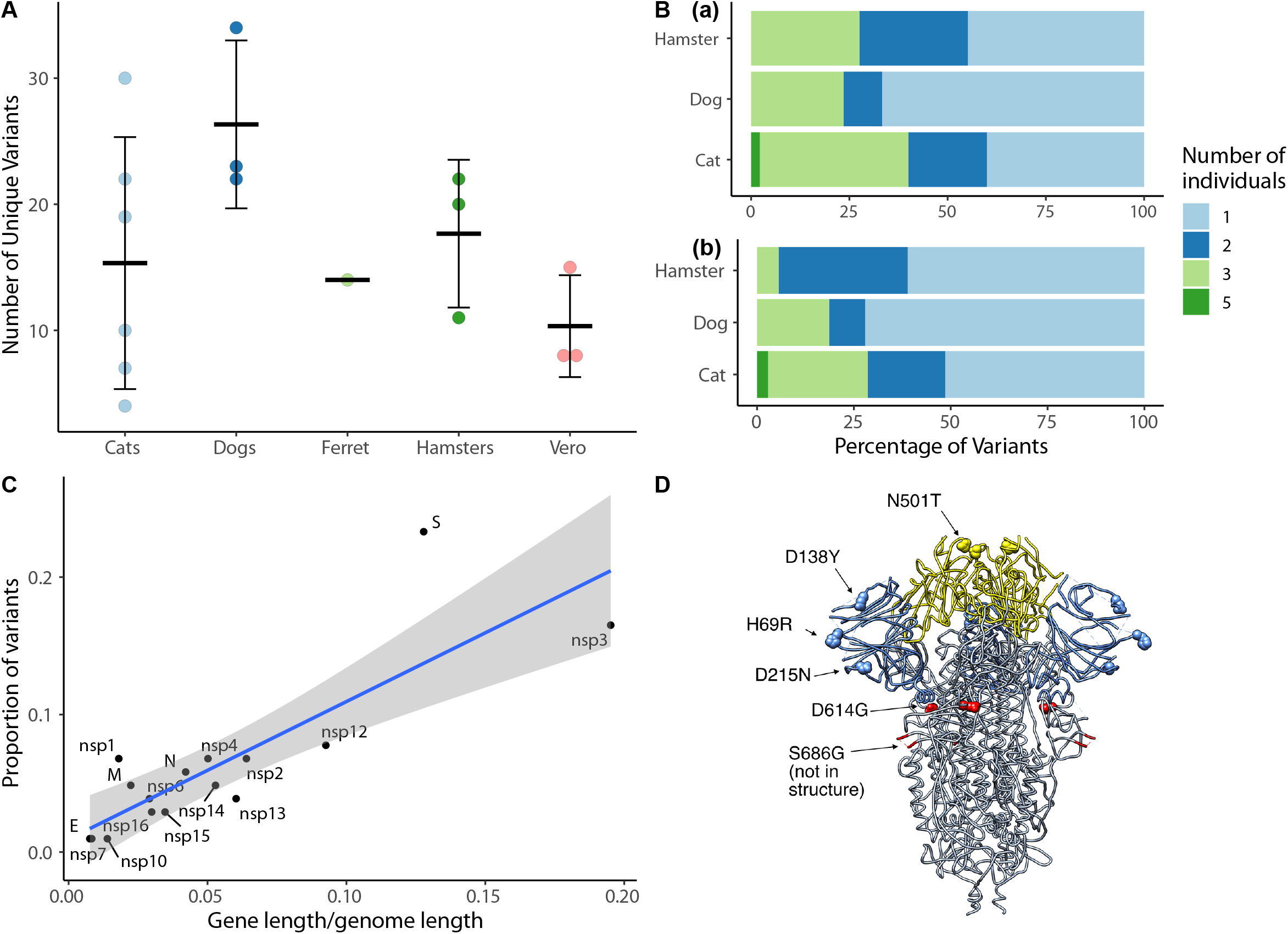
SARS-CoV-2 viral evolution differs across species, gene regions, and individuals. (**A**) Each point indicates the number of unique variants detected at ≥ 3% frequency in SARS-CoV-2 genomes recovered from individual animals. There is no significant difference in the number of unique variants detected in different species (ANOVA, p=0.22). (**B**) Analysis of variant distribution within species reveals that the majority of variants were detected in just one individual within each species. Subplot (a) shows distribution for all variants, while (b) illustrates only variants not occurring in the P3 inoculum at >3%. (**C**) Variants are distributed across viral genes in relation to each gene’s length as a proportion of the entire genome length (linear model, R2=0.69, p<0.0001). The spike protein contained a notably higher proportion of all variants in comparison to its share of genome length. Grey shading represents the 95% confidence interval for the slope of the regression line. (**D**) SARS-CoV-2 spike protein variant “residues of concern” are in N terminal domain (NTD), receptor binding domain (RBD) and furin cleavage site. Residues described in text and Table 1 in the SARS-CoV-2 spike trimer are highlighted on structure 6VXX. Blue indicates NTD, Yellow indicates RBD. The furin cleavage site and adjacent residue 686 are in the indicated loop, which was not resolved in this structure.

Variants were unevenly distributed among individuals of a species. When variants detected in the viral inoculum were excluded from the analysis, a large proportion of variants detected in cats, dogs and hamsters were observed in only one individual of the species (**Fig. 3B**). The SARS-CoV-2 spike gene contained a larger proportion of variants relative to its gene length (**Fig. 3C**), and the majority of variant positions corresponding to human variants of concern were in the spike protein (**Table 1, Fig. 3D**).

Six nonsynonymous and one synonymous amino acid variant were found in all dogs and were present in >50% of sequences recovered from two of three dogs (**Table 2**). These seven variants were found in cats and hamsters at lower frequency. A single novel amino acid variant (D138Y) was detected in all three hamsters. Cats 1-5 were experimentally inoculated; Cat 6 was infected through co-housing with Cat 5 one day post-infection (*6*). Cats were inoculated in two cohorts (Cohort 1=1, 2, 3; Cohort 2= 4, 5, 6). Virus recovered from cats in Cohort 1 had significantly more variants than cats in Cohort 2, but variants were at higher frequency (near fixation) in cats in Cohort 2 (P<0.02). No biological or methodological differences existed between the two experimental cohorts of cats to explain this observed variation. Two amino acid variants (H69R and D614G) were found at >98% in Cats 5 and 6, and S686G was found in 99% of sequences from Cat 1. Six of the ten variants detected in Cat 5, including spike variants H69R and D614G, were also detected in Cat 6, which was infected through contact with Cat 5 (table S2). The single unique variant detected in the contact cat caused a frameshift at D124 in nsp15. Depth of coverage for Cat 5 was <40x at this position, so no variants were called. No amino acid variants from SARS-CoV-2 sequences were shared across all six cats.

### Signatures of selection

Nonsynonymous (πN) and synonymous (πS) nucleotide diversity and πN/πS for SARS-CoV-2 populations was evaluated as a measure of within-host diversity and signatures of selection (**Fig. 4**, table S3, table S4). At the population level, πN was greater than πS for 11 out of 13 SARS-CoV-2 samples (paired t-test, p<0.01), indicating positive selection across hosts (**Fig. 4A**). At the species level, selection of SARS-CoV-2 in dogs and hamsters was most significant (**Fig. 4A**). Further, we recorded a significant difference between πN and πS in orf1ab, spike and membrane proteins (p<0.02, **Fig. 4B**), indicating positive selection at the level of these gene products. In particular, spike, πN was significantly greater than πS for viral genomes recovered from all 16 host and cell culture samples (p<0.0001).

**Fig 4.**
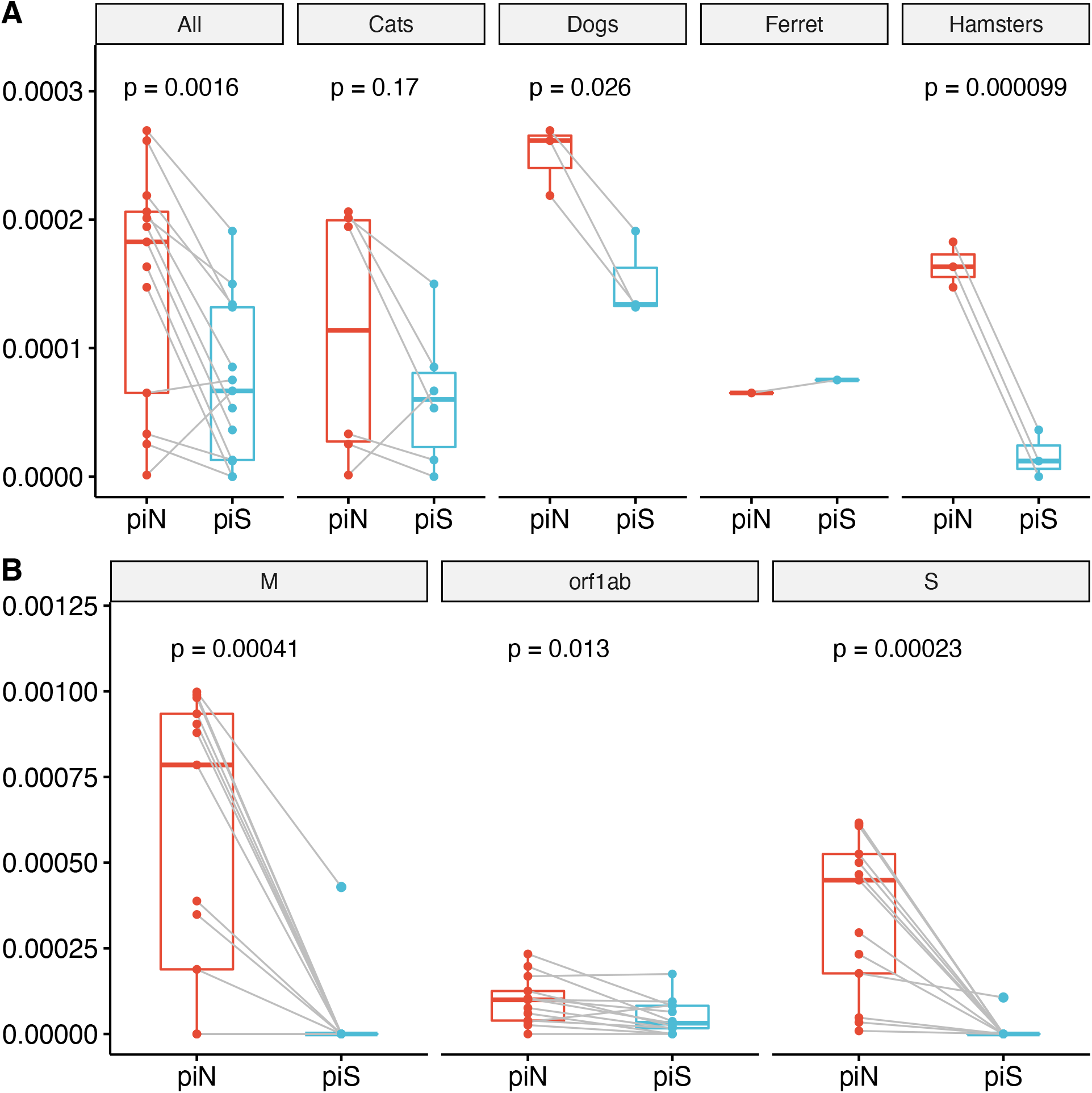
Signatures of positive selection are detected in SARS-CoV-2 genome sequences recovered from experimentally inoculated cats, dogs, hamsters and ferret. (**A**) Comparison of nonsynonymous (πN) and synonymous (πS) nucleotide diversity reveals that πN is significantly greater than πS, indicating positive selection. Each point represents a measurement πN (red) or πS (blue) calculated for the entire SARS-CoV-2 genome from sequences recovered from an individual animal, relative to the reference sequence of USA-WA1/2020. Analysis of the same measures within each species reveals πN > πS for viral genomes is greater in dogs and hamsters. (**B**) Orf1ab, S, and M are undergoing positive selection in mammalian hosts. Each point represents πN or πS calculated for a specific SARS-CoV-2 gene or open reading frame from sequences recovered from an individual animal.

## Discussion

### Cell culture-associated mutation

Variants that reached near fixation in cell culture but reverted to wild type in animal infections suggests a role for these residues in facilitating in vivo versus in vitro infection. Rapid reversion of variants arising in Vero E6 cells (originally derived from African green monkeys) occurred in dogs, cats, and hamsters, suggesting that selective advantages of cell culture variants are due to differences between in vivo and in vitro environments rather than specific host-virus adaptation. For example, in vivo viral attachment and entry in vivo requires circumvention of respiratory epithelial defenses such as mucus and cilia; residues which play roles in overcoming these physical features would not be highly selected in cell culture.

Amino acid R685 is located in the furin cleavage site, a motif unique to SARS-CoV-2 relative to SARS-CoV and other SARS-related coronaviruses (*17*). The PRRAR insertion at this location provides a motif for serine proteases to cleave the spike protein into S1 and S2 subunits, a key step initiating viral entry via membrane fusion. Deletion of the furin cleavage site has been shown to reduce SARS-CoV-2 replication in human respiratory cells, hamster and hACE2 mice (*18*). Previous work has demonstrated rapid emergence of spike variants at the furin-cleavage motif, including R685H, following five passages in type II transmembrane serine protease (TMPRSS2)-deficient cells (*19*). This explains the emergence of this variant in our study following propagation in Vero E6 cells, which do not express high levels of TMPRSS2 (*20*). The distribution of variants arising in vitro (nsp12, spike, N, M; **Fig. 2**) indicates functions beyond receptor binding and entry as drivers of selection, which would be predicted to occur in spike.

### Patterns in viral evolution within and across species

We observed the emergence of low, intermediate and high frequency variants across the SARS-CoV-2 genome in viral RNA recovered from cats, dogs, hamsters and ferrets. Low frequency variants (<25%) may represent mutations occurring during population expansion following infection of a new individual. The majority of variants arising in vivo were in the spike protein, which was under the strongest selective pressure, illustrating the importance of this residue in shaping cross-species transmission events. Variants detected in animals were likely to arise either by rapid selection of low frequency inoculum variants with structural features that favored in vivo attachment, entry, and replication, or as de novo mutations arising from errors during viral replication.

The magnitude of cell culture variant reversion was greater in dogs than other species. All five cell culture-associated variants decreased from >93% to <41%; in particular, D135E and R685H decreased to ≤10%. SARS-CoV-2 was not culturable from canine samples, consistent with previous reports (*21*), and was below the limit of detection by qPCR, though low level seroconversion suggested some viral replication had occurred (*6*). These experimental results contrast with widespread global reports of canine companions becoming infected through contact with their SARS-CoV-2-infected owners (*22-24*). Many variants detected in dogs were not found in the inoculum, reached high frequencies (**Table 2**) and were under strong selective pressure (**Fig. 4**). The majority of canine variants were found in nonstructural replicase genes (**Table 2**), providing strong evidence of selective pressure and host-viral molecular interactions in dogs directed towards selection for viral replication function. The differences between reports of natural reverse-zoonotic infections and laboratory exposures in dogs may stem from disparate doses, dose frequency, or strains (human-adapted or cell culture-propagated virus).

Both domestic and nondomestic cats appear to be highly susceptible to SARS-CoV-2 infection, and readily transmit virus to other cats (*6,7,10,11*). The similarity between sequences recovered from Cat 5 and Cat 6 (exposed via direct contact with Cat 5), demonstrates the ability of novel mutations to be horizontally transmitted (table S2). In contrast to dogs, high viral titers were recovered from cats (*6*), and fewer consensus and shared variants arose in cats than dogs, suggesting that SARS-CoV-2 viral pathogenesis in cats is more similar to humans than dogs. SARS-CoV-2 RNA recovered from hamsters one-day post inoculation had relatively low frequency variants compared to dogs and cats (**Table 2**), but was under strong selective pressure (**Fig. 4**). Virus recovered from one ferret did not demonstrate reversion of Vero E6 cell-passaged variants (**Fig. 2**). However, spike residue 501 was under selection as has been reported in other mustelids (*25,26*).

### Variants of concern

We detected variation at positions that are implicated in the adaptive evolution of SARS-CoV-2 in humans. Some variants, like spike D614G, are identical to variants from human infections. Others, like spike D215N and N501T, represent different substitutions at the same sites as “variants of concern,” which are termed as such due to their potential effects on viral fitness, transmission, replication, or immune evasion. The rapid emergence of these variants in RNA collected 1-3 days post-inoculation illustrates substantial adaptive pressures shaping viral evolution in early stages of host-switching.

Spike N-terminal domain (NTD) variants H69R, D215N, D138Y were detected in cats, dogs and hamsters (**Fig. 3D**). D215N reached highest frequencies in dogs and H69R went to fixation in Cats 5 and 6. In contrast, D215H was present at fixation in the inoculum (**Fig. 2**). In humans the spike mutation D215G is characteristic of the B.1.351 lineage that emerged in South Africa in October 2020 (*27*). Position 215 is also adjacent to an insertion in virus recovered from deer mice (*Peromyscus maniculatus*; (*28*)). D138Y is a defining mutation of the P.1 lineage that emerged in Brazil in December 2020 (*29*). The frequency of NTD variants reported in humans, and in animals in this study, suggests this domain is a hotspot for variants that may have a transmission or replication advantage. At least one neutralizing human antibody that binds to the NTD has been identified (*30*) and this domain has been intensely scrutinized as a vaccine epitope. NTD variants are thus of concern as adaptive changes in this region may facilitate immune evasion.

Amino acid 501, asparagine (N) in original SARS-CoV-2 viruses, is located in the receptor binding domain (RBD) of the spike protein (**Fig. 3D**). This position is a threonine (T) in SARS-CoV and in the closely related coronavirus isolated from a pangolin prior to SARS-CoV-2 spillover (*31*). We observed the rapid emergence of N501T in a ferret. The identical variant arose following experimental infection of ferrets (*25*), and has also emerged repeatedly on mink farms in association with 69-70del (*26*). Thus, these amino acid changes may contribute to adaptation of SARS-CoV-2 to ferret and mink hosts (*32*). The co-occurrence of spike 69-70del and N501Y has been recorded in emergent SARS-CoV-2 lineages in humans (*13,33*). The N501Y substitution has been shown to increase binding affinity to the human ACE2 (*34*).

H655Y, a feature of the Brazilian P.1 lineage in humans, was detected in RNA from three of six cats, all three dogs, and two hamsters, and was present at low frequency (1.1%) in the P3 viral inoculum (**Table 1**). Spike amino acid 655 is located near the S1/S2 cleavage site of the spike protein between the RBD and the fusion peptide. This variant has been reported to be under positive selection in experimentally inoculated cats (*35*) and has been shown to emerge following one passage of pseudotyped SARS-CoV-2 in cell culture in the presence of anti-spike human antibodies (*36*).

Spike variant D614G was the first widely-recognized emergent mutation in human SARS-CoV-2 lineages in early 2020 and has rapidly increased in prevalence to become the dominant sequence worldwide. At the end of February 2021, D614G represented 94.8% SARS-CoV-2 sequences publicly available in the GISAID database (*37*). The growth advantage of this variant is ascribed to replication fitness advantage (*14,38*). We identified D614G in two inoculated cats. This variant was not detectable in the inoculum, and thus represents an example of convergent evolution in humans and cats. The variant reached fixation in Cat 5, and was transmitted to Cat 6 via contact infection, indicating this variant could become established in a cat-to-cat transmission chain.

Variant L37F in nsp6, detected in RNA from cats, dogs and hamsters, also emerged early during the COVID-19 pandemic as a defining mutation of the GISAID Clade V; however, it was not as ultimately successful in spreading in humans as D614G (*37*). We detected L37F resulting from a single nucleotide variation in all four species. Two frameshift mutations (nucleotide insertion and nucleotide deletion) were also detected at L37. Although there is no longer sustained public attention on L37 variants in human SARS CoV-2, previous research has linked the presence of the L37F G>T SNV to asymptomatic infection in humans (*39*).

SARS-CoV-2 is thought to have emerged following passage of a bat-origin virus through an intermediate host, prior to infecting humans (*4,5*). Given the susceptibility of companion animals to SARS-CoV-2 infection and reports of SARS-CoV-2 transmission from humans to animals and then back to humans, the rapid adaptation we document illustrates the potential of reverse zoonosis to accelerate variant emergence for SARS-CoV-2 and other viruses. The rapidity of in vitro and in vivo adaptation underscores the extraordinary plasticity and adaptive potential of SARS-CoV-2. Pathogens are under strong selective pressure to propagate in the host environment, while host defenses are aligned to prevent pathogen replication. This host-pathogen arms race results in varied outcomes that can lead to increases or decreases in virulence and transmission (*40*). Thus, monitoring SARS-CoV-2 evolution in hosts with the potential to infect humans should be a high priority, as de novo novel variants or recombinants between human and animal strains could result in altered transmission pathways and vaccine efficacy (*41,42*).

Our work additionally illustrates that virus evolution and adaptation following passage in cell culture and experimental infection is far more complex than has typically been acknowledged. The advent of new technologies that afford the opportunity to assess viral quasispecies within inocula and biological samples provides an exciting opportunity for future studies of viral evolution and pathogenesis in both humans and animal hosts.

## Materials and Methods

### Cell culture passage in vitro

SARS-CoV-2 strain USA-WA1/2020 (Genbank MN985325.1) was obtained (*15*) and passaged in E6 Vero cells a total of three times. Briefly, 100 μL of viral stock was inoculated onto a flask of confluent E6 Vero cells, allowed to adsorb for 30 minutes, and incubated in cell culture media (Dulbecco’s Modified Eagle Media supplemented with 10% fetal bovine serum and antibiotics) for 3-4 days, harvested and frozen in aliquots.

### Infections in vivo

Intranasal inoculations of cats, dogs, and ferrets with SARS-CoV-2 infection were conducted in ABSL3 facilities at Colorado State University. We conducted a comprehensive analysis of infection outcomes, including assessment of virus neutralization, seroconversion, cat-to-cat transmission, and resistance to reinfection (*6*). SARS-CoV-2 strain USA-WA1/2020 was expanded in Vero E6 cells, and all four species were inoculated with the same “Passage 3” viral stock. Cats 2, 3 and 4 were in Cohort 1, and Cats 1 and 5 were in Cohort 2. Cat 6 was co-housed with Cat 5 for contact transmission of the virus (*6*). Animals were lightly anesthetized and between 10^5^ – 10^6^ pfu SARS-CoV-2 was instilled via the nares. Oropharyngeal and/or nasal samples collected for up to 10 days. Oral swabs were placed in BA-1 medium (Tris-buffered MEM containing 1% BSA) supplemented with gentamicin, amphotericin B and penicillin/streptomycin. Nasal flushes were performed by instilling 1 mL BA-1 dropwise into the nares of awake or lightly anesthetized cats, dogs and ferrets and collecting nasal discharge into a sterile petri dish by allowing the wash fluid to be sneezed out or dripped onto the dish. Viral titers (expressed as plaque-forming units [pfu]) were assessed for all infected individuals (*6*).

### RNA extraction and sequencing

100 μL of lavage fluid was added into 900 μL of trizol to inactivate the virus and prepare the sample for RNA extraction. RNA was extracted using a modified Zymo RNA Clean and Concentrator 5 kit (Zymo Research, Irvine, CA, USA). Samples were thawed on ice and 200 μL of chloroform was added. Samples were agitated by hand for 15 seconds, incubated at room temperature for at least two minutes and centrifuged for 10 minutes at 12,000g. 450 μL of the aqueous phase was removed and placed in 1.5 mL microcentrifuge tubes containing 900 μL of 1:1 Zymo RNA binding buffer and 100% EtOH (450 μL each). Samples were vortexed and then 750 μL of each was transferred to RNA CC-5 columns and centrifuged for 1 minute at 12,000g. Each column was washed with 400 μL of Zymo RNA wash buffer, centrifuged again for 1 minute at 12,000g. A mixture of 24 μL Zymo RNA wash buffer, 3 μL DNase I buffer, and 3 μL DNase I (both New England Biolabs, Ipswich, MA, USA) was added to 30 μL of the mixture and incubated for 22 minutes at room temperature. Following incubation, columns were spun in collection tubes for 30 seconds at 9000g. RNA prep buffer (400 μL) was added to each column and centrifuged again. Flow through was discarded and columns washed with 800 μL RNA wash buffer, followed immediately by a second wash with 400 μL. Wash buffer was discarded and samples were centrifuged for one minute at 16,000g to dry the membrane. We transferred columns to labeled sterile microcentrifuge tubes and to elute RNA. Immediately following RNA extraction cDNA was generated by adding 1 μL of 3µg/μL of random primers (Thermo Fisher, Waltham, MA, USA) and 1 μL of 10 mM dNTPs (New England Biolabs, Ipswich, MA, USA) to 10 μL RNA and incubating at 65°C for 5 min on a C1000 Touch Thermal Cycler (Bio-Rad, USA) followed by chilling on ice. We added 4 μL 5x First-Strand Buffer, 2 μL 0.1M DTT, and 1 μL RNase OUT (40 units/μL) to each sample, pipette mixed, and incubated at 25°C for 2 minutes. 1 μL (200 units) of SuperScript II RT was added pipette mixed, and incubated at 25°C for 10 min, 42°C for 50 min, and 70°C for 15 min. cDNA was frozen at -20°C until use in library preparation.

### Amplicon generation and library preparation

We employed an amplicon based next generation sequencing approach using the primers and protocols developed and optimized for SARS-CoV-2 by the ARTIC Network (*45,46*). All samples were sequenced with ARTIC version 3 primers except for Cat 5 and one replicate of Cat 4, which were sequenced with the previous version 2 primers. This sequencing method generates high coverage of viral genomes from even low template samples by utilizing an initial PCR step resulting in ∼400bp amplicons and primers that span the coding region (*46*). Briefly, two primer pools were used for the initial PCR amplification and then pooled prior to library preparation. We then visualized PCR products on agarose gels and quantified them with the Qubit Broad Range kit (Thermo Fisher, Waltham, MA, USA) and normalized samples to 540ng of cDNA. We prepared sequencing libraries using the New England Biolabs Ultra II kit with the only modification being the use of Ampure XP beads for cleanup and size selection (Beckman Coulter, Brea, CA, USA). Following adapter ligation a single size selection was conducted at 0.65X. We then pooled libraries for sequencing on Illumina MiSeq (Illumina, San Diego, CA, USA) using the v2 500 cycle 2 by 250bp kit.

### Bioinformatics & data analysis

Raw sequencing data were input into a comprehensive Nextflow pipeline for processing next generation sequencing data and single nucleotide and structural variant calling. Nextflow is a bioinformatics workflow manager that facilitates the development of complex and reproducible computational pipelines (*47*). Briefly, data were trimmed for adapters and low quality using Cutadapt (*48*), followed by aligning reads to the viral reference sequence. Data were pre-processed for quality with GATK (*49*) prior to calling single nucleotide and structural variants with LoFreq (*50*). SnpEff and SnpSift were used to annotate variants and predict their functional effects (*51,52*). We designed our variant calling pipeline to ensure that we could differentiate between variants that were not detected due to their absence or presence at a frequency below our detection threshold as compared to inadequacy of coverage depth. The outputs of these analyses were tabulated, processed and visualized in R.

UCSF Chimera software was used to visualize the location of variants in the three-dimensional structure of the SARS-CoV-2 spike protein (*53*). SNPGenie was used to calculate nonsynonymous and synonymous nucleotide diversity (*54*). SNPGenie takes a population of sequences and estimates the mean number of pairwise differences per site, or the nucleotide diversity. Estimates are weighted by allele frequencies. The SNPGenie snpgenie.pl takes as input variant call information contained in .vcf files, a reference genome and genome annotations, and outputs estimates of nucleotide diversity at nonsynonymous and synonymous coding sites at both a genome/population level and by gene product. One advantage of using the nucleotide diversity statistic (π) to compare intrahost viral diversity is that it is not biased by variations in sequencing depth (*55*).

## Acknowledgments

We are grateful to K. Holmes for her thoughtful editorial review and enthusiastic discussions about coronavirus biology. We thank M. Nehring for laboratory management, A. Hartwig and S. Porter for assisting with animal studies, and G. Ebel and N. Sexton for contributing primers. The following reagent was deposited by the Centers for Disease Control and Prevention and obtained through BEI Resources, National Institute of Allergy and Infectious Diseases, NIH: SARS-Related Coronavirus 2, Isolate USA-WA1/2020, NR-52281.

## Author contributions

Conceptualization: SV, MS, RG

Methodology: ABL, MS, RG, LB, SV, RB

Formal Analysis: RG, MS, LB

Visualization: RG, MS, LB

Funding acquisition: SV, RG, MS

Resources: ABL, RB, MS, SV

Supervision: SV

Writing – original draft: RG, SV, LB

Writing – review & editing: ABL, RB, SV, RG, LB, MS

## Data and materials availability

All SARS-CoV-2 raw sequence data used in this study will be publicly available upon publication in the NCBI SRA database under BioProject PRJNA704947. The bioinformatics pipeline used to analyze the data is publicly available at https://github.com/stenglein-lab/viral_variant_caller. The code used to create the three-dimensional visualization of the spike protein is available at https://github.com/stenglein-lab/highlight_residues_on_spike_structure. Additional data including variant summary tables output by the pipeline and R scripts for data processing and visualization are available at https://github.com/laurabashor/SARS-CoV-2-evolution.

## References

1. D. Sauter, F. Kirchhoff, Key Viral Adaptations Preceding the AIDS Pandemic. Cell Host and Microbe 25, 27–38 (2019). doi:10.1016/j.chom.2018.12.002

2. K. G. Andersen, A. Rambaut, W. I. Lipkin, E. C. Holmes, R. F. Garry, The proximal origin of SARS-CoV-2. Nature Medicine 26, 450–452 (2020). doi:10.1038/s41591-020-0820-9

3. P. C. Cross, D. J. Prosser, A. M. Ramey, E. M. Hanks, K. M. Pepin, Confronting models with data: The challenges of estimating disease spillover. Philosophical Transactions of the Royal Society B: Biological Sciences 374, 1782 (2019). doi:10.1098/rstb.2018.0435

4. P. Zhou, X. lou Yang, X. G. Wang, B. Hu, L. Zhang, W. Zhang, H. R. Si, Y. Zhu, B. Li, C. L. Huang, H. D. Chen, J. Chen, Y. Luo, H. Guo, R. di Jiang, M. Q. Liu, Y. Chen, X. R. Shen, X. Wang, X. S. Zheng, K. Zhao, Q. J. Chen, F. Deng, L. L. Liu, B. Yan, F. X. Zhan, Y. Y. Wang, G. F. Xiao, Z. L. Shi, A pneumonia outbreak associated with a new coronavirus of probable bat origin. Nature 579, 270–273 (2020). doi:10.1038/s41586-020-2012-7

5. T. T. Y. Lam, N. Jia, Y. W. Zhang, M. H. H. Shum, J. F. Jiang, H. C. Zhu, Y. G. Tong, Y. X. Shi, X. B. Ni, Y. S. Liao, W. J. Li, B. G. Jiang, W. Wei, T. T. Yuan, K. Zheng, X. M. Cui, J. Li, G. Q. Pei, X. Qiang, W. Y. M. Cheung, L. F. Li, F. F. Sun, S. Qin, J. C. Huang, G. M. Leung, E. C. Holmes, Y. L. Hu, Y. Guan, W. C. Cao, Identifying SARS-CoV-2-related coronaviruses in Malayan pangolins. Nature 583, 282–285 (2020). doi:10.1038/s41586-020-2169-0

6. A. M. Bosco-Lauth, A. E. Hartwig, S. M. Porter, P. W. Gordy, M. Nehring, A. D. Byas, S. VandeWoude, I. K. Ragan, R. M. Maison, R. A. Bowen, Experimental infection of domestic dogs and cats with SARS-CoV-2: Pathogenesis, transmission, and response to reexposure in cats. Proceedings of the National Academy of Sciences of the United States of America 117, 26382–26388 (2020). doi:10.1073/pnas.2013102117

7. D. McAloose, M. Laverack, L. Wang, M. L. Killian, L. C. Caserta, F. Yuan, P. K. Mitchell, K. Queen, M. R. Mauldin, B. D. Cronk, S. L. Bartlett, J. M. Sykes, S. Zec, T. Stokol, K. Ingerman, M. A. Delaney, R. Fredrickson, M. Ivančić, M. Jenkins-Moore, K. Mozingo, K. Franzen, N. H. Bergeson, L. Goodman, H. Wang, Y. Fang, C. Olmstead, C. McCann, P. Thomas, E. Goodrich, F. Elvinger, D. C. Smith, S. Tong, S. Slavinski, P. P. Calle, K. Terio, M. K. Torchetti, D. G. Diel, From people to panthera: Natural sars-cov-2 infection in tigers and lions at the bronx zoo. mBio 11, 1–13 (2020). doi:10.1128/mBio.02220-20

8. B. B. O. Munnink, R. S. Sikkema, D. F. Nieuwenhuijse, R. J. Molenaar, E. Munger, R. Molenkamp, A. van der Spek, P. Tolsma, A. Rietveld, M. Brouwer, N. Bouwmeester-Vincken, F. Harders, R. H. van der Honing, M. C. A. Wegdam-Blans, R. J. Bouwstra, C. GeurtsvanKessel, A. A. van der Eijk, F. C. Velkers, L. A. M. Smit, A. Stegeman, W. H. M. van der Poel, M. P. G. Koopmans, Transmission of SARS-CoV-2 on mink farms between humans and mink and back to humans. Science 371, 172–177 (2021). doi:10.1126/science.abe5901

9. M. A. Shereen, S. Khan, A. Kazmi, N. Bashir, R. Siddique, COVID-19 infection: Origin, transmission, and characteristics of human coronaviruses. Journal of Advanced Research 24, 91–98 (2020). doi:10.1016/j.jare.2020.03.005

10. A. Banerjee, K. Mossman, M. L. Baker, Zooanthroponotic potential of SARS-CoV-2 and implications of re-introduction into human population. Cell Host & Microbe 29, 160–164 (2021). doi:10.1016/j.chom.2021.01.004

11. H. D. Hedman, E. Krawczyk, Y. A. Helmy, L. Zhang, C. Varga, L. Gralinski, L. Martinez-Sobrido, pathogens Host Diversity and Potential Transmission Pathways of SARS-CoV-2 at the Human-Animal Interface. Pathogens 10, 180 (2021). doi:10.3390/pathogens10020180

12. F. Wu, S. Zhao, B. Yu, Y. M. Chen, W. Wang, Z. G. Song, Y. Hu, Z. W. Tao, J. H. Tian, Y. Y. Pei, M. L. Yuan, Y. L. Zhang, F. H. Dai, Y. Liu, Q. M. Wang, J. J. Zheng, L. Xu, E. C. Holmes, Y. Z. Zhang, A new coronavirus associated with human respiratory disease in China. Nature 579, 265–269 (2020). doi:10.1038/s41586-020-2008-3

13. A. Rambaut, N. Loman, O. Pybus, W. Barclay, J. Barrett, A. Carabelli, T. Connor, T. Peacock, D. L. Robertson, E. Volz, Preliminary genomic characterisation of an emergent SARS-CoV-2 lineage in the UK defined by a novel set of spike mutations (Virological, 2020); https://virological.org/t/preliminary-genomic-characterisation-of-an-emergent-sars-cov-2-lineage-in-the-uk-defined-by-a-novel-set-of-spike-mutations/563.

14. B. Korber, W. M. Fischer, S. Gnanakaran, H. Yoon, J. Theiler, W. Abfalterer, N. Hengartner, E. E. Giorgi, T. Bhattacharya, B. Foley, K. M. Hastie, M. D. Parker, D. G. Partridge, C. M. Evans, T. M. Freeman, T. I. de Silva, A. Angyal, R. L. Brown, L. Carrilero, L. R. Green, D. C. Groves, K. J. Johnson, A. J. Keeley, B. B. Lindsey, P. J. Parsons, M. Raza, S. Rowland-Jones, N. Smith, R. M. Tucker, D. Wang, M. D. Wyles, C. McDanal, L. G. Perez, H. Tang, A. Moon-Walker, S. P. Whelan, C. C. LaBranche, E. O. Saphire, D. C. Montefiori, Tracking Changes in SARS-CoV-2 Spike: Evidence that D614G Increases Infectivity of the COVID-19 Virus. Cell 182, 812–827 (2020). doi:10.1016/j.cell.2020.06.043

15. J. Harcourt, A. Tamin, X. Lu, S. Kamili, S. K. Sakthivel, J. Murray, K. Queen, Y. Tao, C. R. Paden, J. Zhang, Y. Li, A. Uehara, H. Wang, C. Goldsmith, H. A. Bullock, L. Wang, B. Whitaker, B. Lynch, R. Gautam, C. Schindewolf, K. G. Lokugamage, D. Scharton, J. A. Plante, D. Mirchandani, S. G. Widen, K. Narayanan, S. Makino, T. G. Ksiazek, K. S. Plante, S. C. Weaver, S. Lindstrom, S. Tong, V. D. Menachery, N. J. Thornburg, Severe acute respiratory syndrome coronavirus 2 from patient with coronavirus disease, United States. Emerging Infectious Diseases 26, 1266–1273 (2020). doi:10.3201/EID2606.200516

16. N. D. Grubaugh, K. Gangavarapu, J. Quick, N. L. Matteson, J. G. de Jesus, B. J. Main, A. L. Tan, L. M. Paul, D. E. Brackney, S. Grewal, N. Gurfield, K. K. A. van Rompay, S. Isern, S. F. Michael, L. L. Coffey, N. J. Loman, K. G. Andersen, An amplicon-based sequencing framework for accurately measuring intrahost virus diversity using PrimalSeq and iVar. Genome Biology 20, 8 (2019). doi:10.1186/s13059-018-1618-7

17. B. Coutard, C. Valle, X. de Lamballerie, B. Canard, N. G. Seidah, E. Decroly, The spike glycoprotein of the new coronavirus 2019-nCoV contains a furin-like cleavage site absent in CoV of the same clade. Antiviral Research 176, 104742 (2020). doi:10.1016/j.antiviral.2020.104742

18. B. A. Johnson, X. Xie, A. L. Bailey, B. Kalveram, K. G. Lokugamage, A. Muruato, J. Zou, X. Zhang, T. Juelich, J. K. Smith, L. Zhang, N. Bopp, C. Schindewolf, M. Vu, A. Vanderheiden, E. S. Winkler, D. Swetnam, J. A. Plante, P. Aguilar, K. S. Plante, V. Popov, B. Lee, S. C. Weaver, M. S. Suthar, A. L. Routh, P. Ren, Z. Ku, Z. An, K. Debbink, M. S. Diamond, P. Y. Shi, A. N. Freiberg, V. D. Menachery, Loss of furin cleavage site attenuates SARS-CoV-2 pathogenesis. Nature 1–7 (2021). doi:10.1038/s41586-021-03237-4

19. M. Sasaki, K. Uemura, A. Sato, S. Toba, T. Sanaki, K. Maenaka, W. W. Hall, Y. Orba, H. Sawa, SARS-CoV-2 variants with mutations at the S1/S2 cleavage site are generated in vitro during propagation in TMPRSS2-deficient cells. PLOS Pathogens 17, e1009233 (2021). doi:10.1371/journal.ppat.1009233

20. T. Ou, H. Mou, L. Zhang, A. Ojha, H. Choe, M. Farzan, Hydroxychloroquine-mediated inhibition of SARS-CoV-2 entry is attenuated by TMPRSS2. PLOS Pathogens 17, e1009212 (2021). doi:10.1371/journal.ppat.1009212

21. J. Shi, Z. Wen, G. Zhong, H. Yang, C. Wang, B. Huang, R. Liu, X. He, L. Shuai, Z. Sun, Y. Zhao, P. Liu, L. Liang, P. Cui, J. Wang, X. Zhang, Y. Guan, W. Tan, G. Wu, H. Chen, Z. Bu, Z. Bu, Susceptibility of ferrets, cats, dogs, and other domesticated animals to SARS-coronavirus 2. Science 368, 1016–1020 (2020). doi:10.1126/science.abb7015

22. T. H. C. Sit, C. J. Brackman, S. M. Ip, K. W. S. Tam, P. Y. T. Law, E. M. W. To, V. Y. T. Yu, L. D. Sims, D. N. C. Tsang, D. K. W. Chu, R. A. P. M. Perera, L. L. M. Poon, M. Peiris, Infection of dogs with SARS-CoV-2. Nature 586, 776–778 (2020). doi:10.1038/s41586-020-2334-5

23. A. J. Perisé-Barrios, B. D. Tomeo-Martín, P. Gómez-Ochoa, P. Delgado-Bonet, P. Plaza, P. Palau-Concejo, J. González, G. Ortiz-Díez, A. Meléndez-Lazo, M. Gentil, J. García-Castro Barbero-Fernández, Humoral responses to SARS-CoV-2 by healthy and sick dogs during the COVID-19 pandemic in Spain. Veterinary Research 52, 22 (2021).

24. E. I. Patterson, G. Elia, A. Grassi, A. Giordano, C. Desario, M. Medardo, S. L. Smith, E. R. Anderson, T. Prince, G. T. Patterson, E. Lorusso, M. S. Lucente, G. Lanave, S. Lauzi, U. Bonfanti, A. Stranieri, V. Martella, F. Solari Basano, V. R. Barrs, A. D. Radford, U. Agrimi, G. L. Hughes, S. Paltrinieri, N. Decaro, Evidence of exposure to SARS-CoV-2 in cats and dogs from households in Italy. Nature Communications 11, 1–5 (2020). doi:10.1038/s41467-020-20097-0

25. M. Richard, A. Kok, D. de Meulder, T. M. Bestebroer, M. M. Lamers, N. M. A. Okba, M. Fentener van Vlissingen, B. Rockx, B. L. Haagmans, M. P. G. Koopmans, R. A. M. Fouchier, S. Herfst, SARS-CoV-2 is transmitted via contact and via the air between ferrets. Nature Communications 11, 1–6 (2020). doi:10.1038/s41467-020-17367-2

26. L. van Dorp, C. C. S. Tan, S. D. Lam, D. Richard, C. Owen, D. Berchtold, C. Orengo, F. Balloux, Recurrent mutations in SARS-CoV-2 genomes isolated from mink point to rapid host-adaptation. bioRxiv 2020.11.16.384743 [Preprint]. 16 November 2020. https://doi.org/10.1101/2020.11.16.384743.

27. H. Tegally, E. Wilkinson, M. Giovanetti, A. Iranzadeh, V. Fonseca, J. Giandhari, D. Doolabh, S. Pillay, E. J. San, N. Msomi, K. Mlisana, A. von Gottberg, S. Walaza, M. Allam Ismail, T. Mohale, A. J. Glass, S. Engelbrecht, G. van Zyl, W. Preiser, F. Petruccione, A. Sigal, D. Hardie, G. Marais, M. Hsiao, S. Korsman, M. A. Davies, L. Tyers, I. Mudau, D. York, C. Maslo, D. Goedhals, S. Abrahams, O. Laguda-Akingba, A. Alisoltani-Dehkordi, A. Godzik, C. K. Wibmer, B. T. Sewell, J. Lourenço, L. C. J. Alcantara, S. L. Kosakovsky Pond, S. Weaver, D. Martin, R. J. Lessells, J. N. Bhiman, C. Williamson, T. de Oliveira, Emergence and rapid spread of a new severe acute respiratory syndrome-related coronavirus 2 (SARS-CoV-2) lineage with multiple spike mutations in South Africa. medRxiv 2020.12.21.20248640 [Preprint]. 22 December 2020. https://doi.org/10.1101/2020.12.21.20248640.

28. A. Fagre, J. Lewis, M. Eckley, S. Zhan, S. M. Rocha, N. R. Sexton, B. Burke, B. Geiss, O. Peersen, R. Kading, J. Rovnak, G. D. Ebel, R. B. Tjalkens, T. Aboellail, T. Schountz, SARS-CoV-2 infection, neuropathogenesis and transmission among deer mice: Implications for reverse zoonosis to New World rodents. bioRxiv 2020.08.07.241810 [Preprint]. 7 August 2020. https://doi.org/10.1101/2020.08.07.241810.

29. N. R. Faria, I. M. Morales Claro, D. Candido, L. A. Moyses Franco, P. S. Andrade, T. M. Coletti, C. A. M. Silva, F. C. Sales, E. R. Manuli, R. S. Aguiar, N. Gaburo, C. da C. Camilo, N. A. Fraiji, M. A. Esashika Crispim, M. do Perpétuo S. S. Carvalho, A. Rambaut, N. Loman, O. G. Pybus, E. C. Sabino, Genomic characterization of an emergent SARS-CoV-2 lineage in Manaus: preliminary findings (Virological, 2020); https://virological.org/t/genomic-characterisation-of-an-emergent-sars-cov-2-lineage-in-manaus-preliminary-findings/586.

30. X. Chi, R. Yan, J. Zhang, G. Zhang, Y. Zhang, M. Hao, Z. Zhang, P. Fan, Y. Dong, Y. Yang, Z. Chen, Y. Guo, J. Zhang, Y. Li, X. Song, Y. Chen, L. Xia, L. Fu, L. Hou, J. Xu, C. Yu, J. Li, Q. Zhou, W. Chen, A neutralizing human antibody binds to the N-terminal domain of the Spike protein of SARS-CoV-2. Science 369, 650–655 (2020). doi:10.1126/science.abc6952

31. X. Tang, C. Wu, X. Li, Y. Song, X. Yao, X. Wu, Y. Duan, H. Zhang, Y. Wang, Z. Qian, J. Cui, J. Lu, On the origin and continuing evolution of SARS-CoV-2. National Science Review 7, 1012–1023 (2020). doi:10.1093/nsr/nwaa036

32. M. R. A. Welkers, A. X. Han, C. B. E. M. Reusken, D. Eggink, Possible host-adaptation of SARS-CoV-2 due to improved ACE2 receptor binding in mink. Virus Evolution 7 (2021). doi:10.1093/ve/veaa094

33. S. A. Kemp, R. P. Datir, D. A. Collier, I. A. T. M. Ferreira, A. Carabelli, W. Harvey, D. L. Robertson, R. K. Gupta, Recurrent emergence and transmission of a SARS-CoV-2 Spike deletion ∆H69/V70. bioRxiv 2020.12.14.422555 [Preprint]. 9 February 2021. https://doi.org/10.1101/2020.12.14.422555.

34. K. K. Chan, T. J. C. Tan, K. K. Narayanan, E. Procko, An engineered decoy receptor for SARS-CoV-2 broadly binds protein S sequence variants. bioRxiv 2020.10.18.344622 [Preprint]. 20 December 2020. https://doi.org/10.1101/2020.10.18.344622.

35. K. M. Braun, G. K. Moreno, P. J. Halfmann, D. A. Baker, E. C. Boehm, A. M. Weiler, A. K. Haj, M. Hatta, S. Chiba, T. Maemura, Y. Kawaoka, K. Koelle, D. H. O’Connor, T. C. Friedrich, Transmission of SARS-CoV-2 in domestic cats imposes a narrow bottleneck. bioRxiv 2020.11.16.384917 [Preprint]. 4 January 2021. https://doi.org/10.1101/2020.11.16.384917.

36. A. Baum, B. O. Fulton, E. Wloga, R. Copin, K. E. Pascal, V. Russo, S. Giordano, K. Lanza, N. Negron, M. Ni, Y. Wei, G. S. Atwal, A. J. Murphy, N. Stahl, G. D. Yancopoulos, C. A. Kyratsous, Antibody cocktail to SARS-CoV-2 spike protein prevents rapid mutational escape seen with individual antibodies. Science 369, 1014–1018 (2020). doi:10.1126/science.abd0831

37. Y. Shu, J. McCauley, GISAID: Global initiative on sharing all influenza data – from vision to reality. Eurosurveillance 22, 30494 (2017). doi:10.2807/1560-7917.ES.2017.22.13.30494

38. N. D. Grubaugh, W. P. Hanage, A. L. Rasmussen, Making Sense of Mutation: What D614G Means for the COVID-19 Pandemic Remains Unclear. Cell 182, 794–795 (2020). doi:10.1016/j.cell.2020.06.040

39. R. Wang, J. Chen, Y. Hozumi, C. Yin, G. W. Wei, Decoding Asymptomatic COVID-19 Infection and Transmission. Journal of Physical Chemistry Letters 11, 10007–10015 (2020). doi:10.1021/acs.jpclett.0c02765

40. B. Longdon, J. D. Hadfield, J. P. Day, S. C. L. Smith, J. E. McGonigle, R. Cogni, C. Cao, F. M. Jiggins, The Causes and Consequences of Changes in Virulence following Pathogen Host Shifts. PLOS Pathogens. 11, e1004728 (2015). doi:10.1371/journal.ppat.1004728

41. Z. Wang, F. Schmidt, Y. Weisblum, F. Muecksch, S. Finkin, D. Schaefer-Babajew, M. Cipolla, C. Gaebler, J. A. Lieberman, T. Y. Oliveira, Z. Yang, M. E. Abernathy, A. Hurley, M. Turroja, K. A. West, K. Gordon, G. Millard, V. Ramos, J. da Silva, J. Xu, R. A. Colbert, R. Patel, J. Dizon, C. Unson-O, I. Shimeliovich, A. Gazumyan, M. Caskey, P. J. Bjorkman, R. Casellas, T. Hatziioannou, P. D. Bieniasz, M. C. Nussenzweig, mRNA vaccine-elicited antibodies to SARS-CoV-2 and circulating variants. biorxiv 2021.01.15.426911 [Preprint]. 30 January 2021. https://doi.org/10.1101/2021.01.15.426911.

42. Novavax Inc., “Novavax COVID-19 Vaccine Demonstrates 89.3% Efficacy in UK Phase 3 Trial” (Novavax, 2021); https://ir.novavax.com/news-releases/news-release-details/novavax-covid-19-vaccine-demonstrates-893-efficacy-uk-phase-3.

43. H. Gu, Q. Chen, G. Yang, L. He, H. Fan, Y. Q. Deng, Y. Wang, Y. Teng, Z. Zhao, Y. Cui, Y. Li, X. F. Li, J. Li, N. N. Zhang, X. Yang, S. Chen, Y. Guo, G. Zhao, X. Wang, D. Y. Luo, H. Wang, X. Yang, Y. Li, G. Han, Y. He, X. Zhou, S. Geng, X. Sheng, S. Jiang, S. Sun, C. F. Qin, Y. Zhou, Adaptation of SARS-CoV-2 in BALB/c mice for testing vaccine efficacy. Science 369, 1603–1607 (2020). doi:10.1126/science.abc4730

44. J. F. W. Chan, A. J. Zhang, S. Yuan, V. K. M. Poon, C. C. S. Chan, A. C. Y. Lee, W. M. Chan, Z. Fan, H. W. Tsoi, L. Wen, R. Liang, J. Cao, Y. Chen, K. Tang, C. Luo, J. P. Cai, K. H. Kok, H. Chu, K. H. Chan, S. Sridhar, Z. Chen, H. Chen, K. K. W. To, K. Y. Yuen, Simulation of the Clinical and Pathological Manifestations of Coronavirus Disease 2019 (COVID-19) in a Golden Syrian Hamster Model: Implications for Disease Pathogenesis and Transmissibility. Clinical Infectious Diseases 71, 2428–2446 (2020). doi:10.1093/cid/ciaa325

45. K. Itokawa, T. Sekizuka, M. Hashino, R. Tanaka, M. Kuroda, Disentangling primer interactions improves SARS-CoV-2 genome sequencing by multiplex tiling PCR. PLOS ONE 15, e0239403 (2020). doi:10.1371/journal.pone.0239403

46. J. Quick, N. D. Grubaugh, S. T. Pullan, I. M. Claro, A. D. Smith, K. Gangavarapu, G. Oliveira, R. Robles-Sikisaka, T. F. Rogers, N. A. Beutler, D. R. Burton, L. L. Lewis-Ximenez, J. G. de Jesus, M. Giovanetti, S. C. Hill, A. Black, T. Bedford, M. W. Carroll, M. Nunes, L. C. Alcantara, E. C. Sabino, S. A. Baylis, N. R. Faria, M. Loose, J. T. Simpson, O. G. Pybus, K. G. Andersen, N. J. Loman, Multiplex PCR method for MinION and Illumina sequencing of Zika and other virus genomes directly from clinical samples. Nature Protocols 12, 1261–1266 (2017). doi:10.1038/nprot.2017.066

47. P. di Tommaso, M. Chatzou, E. W. Floden, P. P. Barja, E. Palumbo, C. Notredame, Nextflow enables reproducible computational workflows. Nature Biotechnology 35 316–319 (2017). doi:10.1038/nbt.3820

48. M. Martin, Cutadapt removes adapter sequences from high-throughput sequencing reads. EMBnet.journal. 17 10 (2011). doi:10.14806/ej.17.1.200

49. A. McKenna, M. Hanna, E. Banks, A. Sivachenko, K. Cibulskis, A. Kernytsky, K. Garimella, D. Altshuler, S. Gabriel, M. Daly, M. A. DePristo, The genome analysis toolkit: A MapReduce framework for analyzing next-generation DNA sequencing data. Genome Research 20, 1297–1303 (2010). doi:10.1101/gr.107524.110

50. A. Wilm, P. P. K. Aw, D. Bertrand, G. H. T. Yeo, S. H. Ong, C. H. Wong, C. C. Khor, R. Petric, M. L. Hibberd, N. Nagarajan, LoFreq: A sequence-quality aware, ultra-sensitive variant caller for uncovering cell-population heterogeneity from high-throughput sequencing datasets. Nucleic Acids Research 40, 11189–11201 (2012). doi:10.1093/nar/gks918

51. P. Cingolani, V. M. Patel, M. Coon, T. Nguyen, S. J. Land, D. M. Ruden, X. Lu, Using Drosophila melanogaster as a model for genotoxic chemical mutational studies with a new program, SnpSift. Frontiers in Genetics 3 (2012). doi:10.3389/fgene.2012.00035

52. P. Cingolani, A. Platts, L. L. Wang, M. Coon, T. Nguyen, L. Wang, S. J. Land, X. Lu, D. M. Ruden, A program for annotating and predicting the effects of single nucleotide polymorphisms, SnpEff: SNPs in the genome of Drosophila melanogaster strain w1118; iso-2; iso-3. Fly 6, 80–92 (2012). doi:10.4161/fly.19695

53. E. F. Pettersen, T. D. Goddard, C. C. Huang, G. S. Couch, D. M. Greenblatt, E. C. Meng, T. E. Ferrin, UCSF Chimera - A visualization system for exploratory research and analysis. Journal of Computational Chemistry 25, 1605–1612 (2004). doi:10.1002/jcc.20084

54. C. W. Nelson, A. L. Hughes, Within-host nucleotide diversity of virus populations: Insights from next-generation sequencing. Infection, Genetics and Evolution 30, 1–7 (2015). doi:10.1016/j.meegid.2014.11.026

55. L. Zhao, C. J. R. Illingworth, Measurements of intrahost viral diversity require an unbiased diversity metric. Virus Evolution 5 (2019). doi:10.1093/ve/vey041

