## Supplementary Materials for "SARS-CoV-2 evolution in animals suggests mechanisms for rapid variant selection"

**This PDF file includes:**

Figs. S1 and S2
Tables S1 to S4

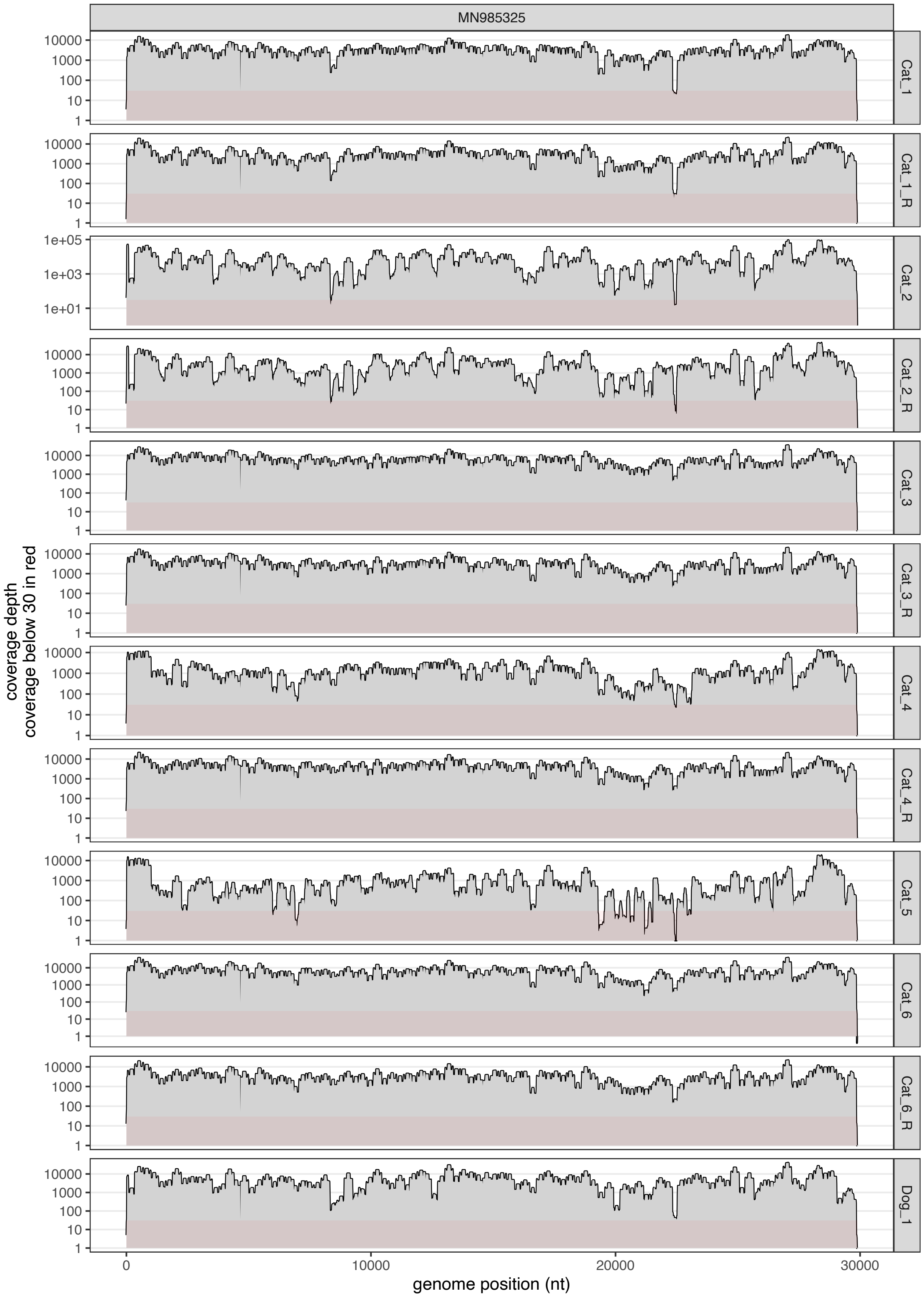
MN985325
Cat_1
Cat_1_R
Cat_2
Cat_2_R
Cat_3
Cat_3_R
Cat_4
Cat_4_R
Cat_5
Cat_6
Cat_6_R
Dog_1
coverage depth
coverage below 30 in red
genome position (nt)
0
10000
20000
30000

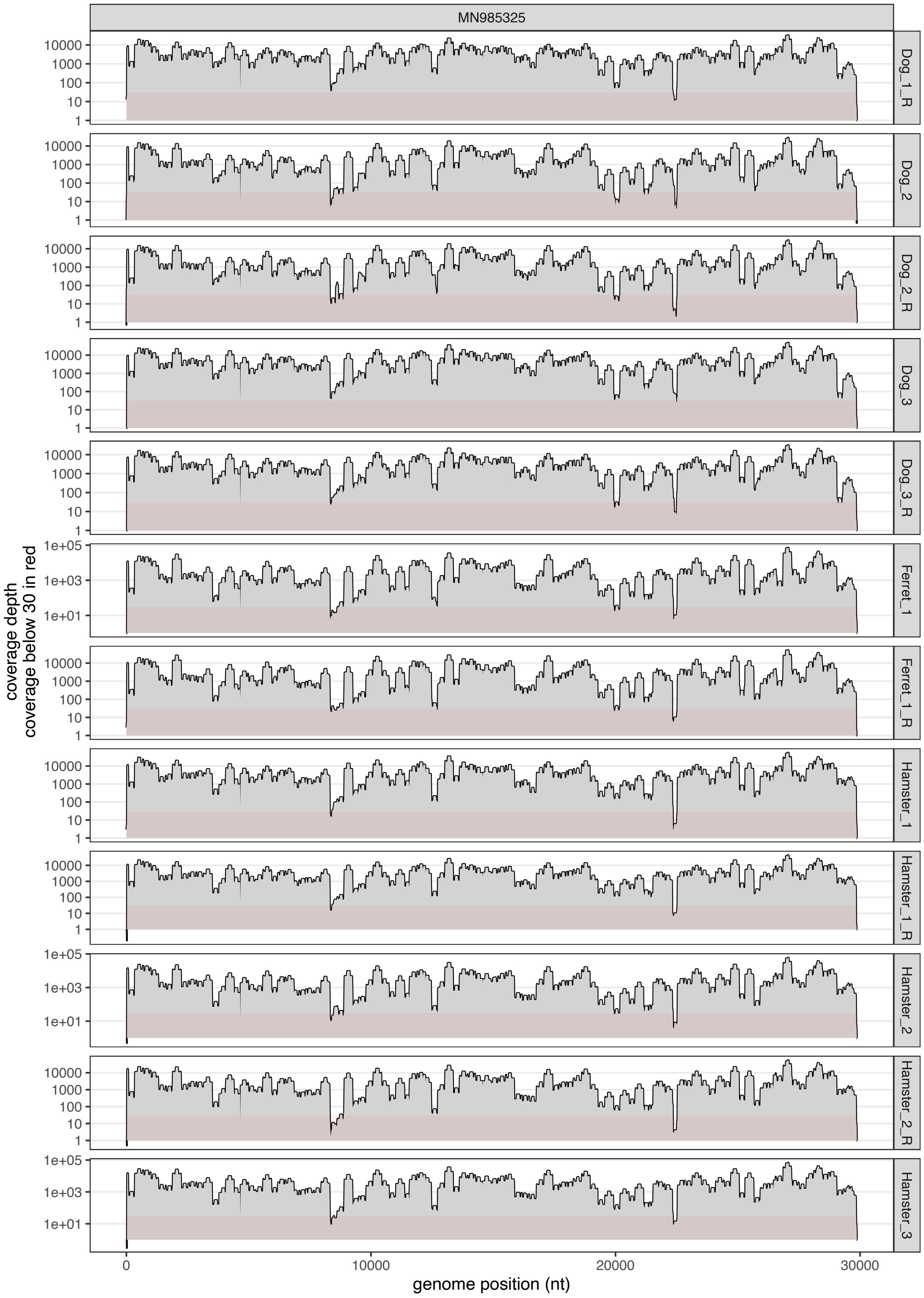
MN985325
coverage depth
coverage below 30 in red
Dog_1_R
Dog_2
Dog_2_R
Dog_3
Dog_3_R
Ferret_1
Ferret_1_R
Hamster_1
Hamster_1_R
Hamster_2
Hamster_2_R
Hamster_3
1e+05
1e+03
1e+01
10000
1000
100
10
1
0
10000
20000
30000
genome position (nt)

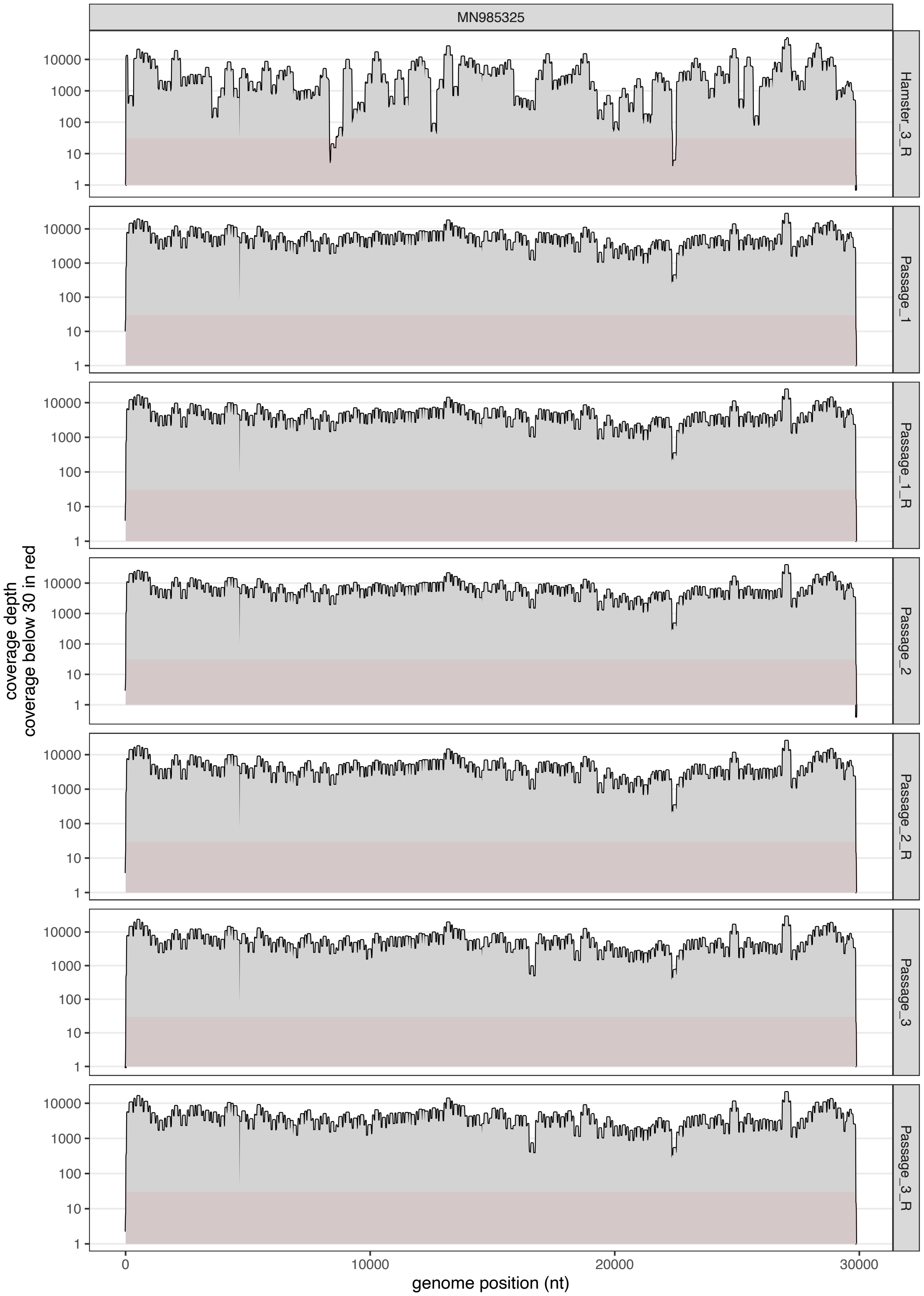


**Fig. S1.**

35 Depth of coverage across the SARS-CoV-2 genome.

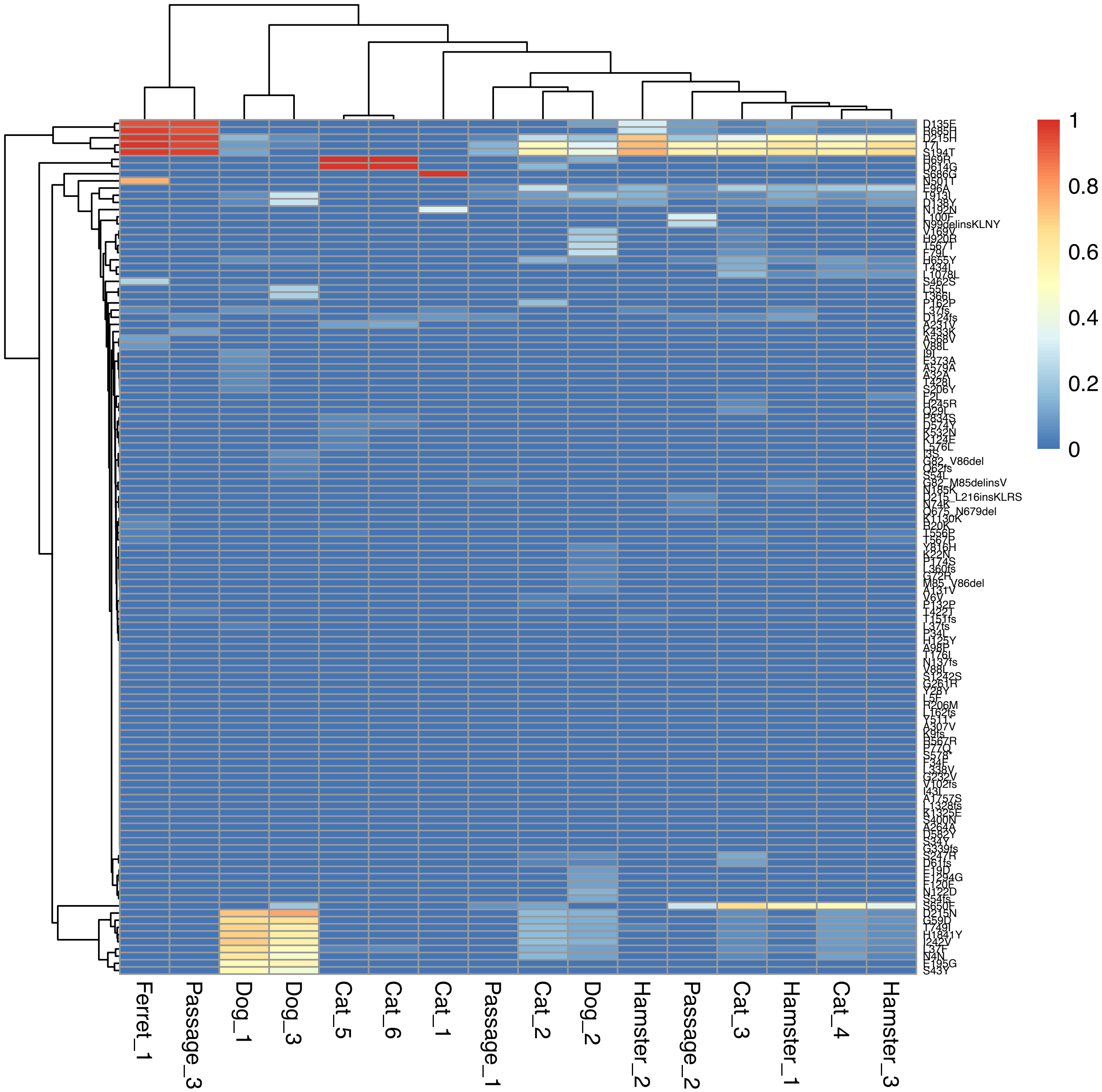


**Fig. S2.**
Heatmap of SARS-CoV-2 variants across samples recovered from cell culture and experimentally inoculated animals.

**Table S1.**
Selected metadata for experimentally inoculated animals included in this study.

| Sample ID | Age | Sex | Inoculation | Sample | log pfu/mL recovered | dpi |
|---|---|---|---|---|---|---|
| Dog 1 | adult (5-8 years) | female | nasal | nasal | <1 | 3 |
| Dog 2 | adult (5-8 years) | female | nasal | nasal | <1 | 3 |
| Dog 3 | adult (5-8 years) | female | nasal | oral | <1 | 3 |
| Ferret 1 | juvenile | | nasal | nasal | | |
| Hamster 1 | juvenile | | nasal | oral | 3.3 | 1 |
| Hamster 2 | juvenile | | nasal | oral | 3.5 | 1 |
| Hamster 3 | juvenile | | nasal | oral | <1 | 1 |
| Cat 1 | adult (5-8 years) | female | nasal | nasal | 6.6 | 3 |
| Cat 2 | adult (5-8 years) | male | nasal | nasal | 5.3 | 3 |
| Cat 3 | adult (5-8 years) | female | nasal | nasal | 6.0 | 3 |
| Cat 4 | adult (5-8 years) | female | nasal | nasal | 5.4 | 3 |
| Cat 5 | adult (5-8 years) | female | nasal | nasal | 4.4 | 3 |
| Cat 6 | adult (5-8 years) | female | contact | nasal | 5.1 | |

**Table S2.**

SARS-CoV-2 variants detected in Cats 5 and 6.

| Variant | Position | CDS | N or S | Cat 5 | Cat 6 |
|---|---|---|---|---|---|
| A231V | 9246 | nsp4 | missense | 0.11 | 0.13 |
| L37F | 11083 | nsp6 | missense | 0.05 | 0.06 |
| K124E | 13394 | nsp10 | missense | 0.04 | 0 |
| K532N | 15036 | nsp12 | missense | 0.06 | 0 |
| T556P | 15106 | nsp12 | missense | 0.03 | 0 |
| L576L | 15168 | nsp12 | synonymous | 0.05 | 0 |
| P834S | 15940 | nsp12 | missense | 0.06 | 0.05 |
| D124fs | 19983 | nsp15 | frameshift variant & stop gained | -- | 0.06 |
| H69R | 21768 | S | missense | 1.0 | 0.99 |
| D574Y | 23282 | S | missense | 0.04 | 0.05 |
| D614G | 23403 | S | missense | 0.99 | 0.98 |

**Table S3.**

Measurements of nucleotide diversity for each sample at the population level and associated statistical analyses.

| Sample ID | pi | piN | piS | piN/piS | p-value | t-value |
|---|---|---|---|---|---|---|
| Cat 1 | 1.60E-05 | 1.19E-06 | 6.67E-05 | 1.78E-02 | | |
| Cat 2 | 1.84E-04 | 2.01E-04 | 1.50E-04 | 1.34E+00 | | |
| Cat 3 | 1.73E-04 | 2.06E-04 | 8.53E-05 | 2.42E+00 | 2.64E-02 | 6.04E+00 |
| Cat 4 | 1.57E-04 | 1.95E-04 | 5.33E-05 | 3.65E+00 | | |
| Cat 5 | 2.77E-05 | 3.32E-05 | 1.29E-05 | 2.57E+00 | | |
| Cat 6 | 1.89E-05 | 2.53E-05 | 0.00E+00 | | | |
| Dog 1 | 1.94E-04 | 2.19E-04 | 1.34E-04 | 1.63E+00 | | |
| Dog 2 | 2.45E-04 | 2.69E-04 | 1.91E-04 | 1.41E+00 | 1.71E-01 | 1.60E+00 |
| Dog 3 | 2.25E-04 | 2.62E-04 | 1.32E-04 | 1.98E+00 | | |
| Ferret 1 | 6.57E-05 | 6.51E-05 | 7.52E-05 | 8.66E-01 | | |
| Hamster 1 | 1.25E-04 | 1.63E-04 | 1.21E-05 | 1.35E+01 | | |
| Hamster 2 | 1.10E-04 | 1.47E-04 | 0.00E+00 | | 9.90E-05 | 1.01E+02 |
| Hamster 3 | 1.45E-04 | 1.83E-04 | 3.63E-05 | 5.03E+00 | | |

**Table S4.**

Measurements of nucleotide diversity for each sample at the gene product level and associated statistical analyses.

| Sample ID | Gene product | piN | piS | piN/piS | p-value | t-value |
|---|---|---|---|---|---|---|
| Cat 1 | E | 0.00E+00 | 0.00E+00 | | | |
| Cat 2 | E | 0.00E+00 | 0.00E+00 | | | |
| Cat 3 | E | 0.00E+00 | 0.00E+00 | | | |
| Cat 4 | E | 0.00E+00 | 0.00E+00 | | | |
| Cat 5 | E | 0.00E+00 | 0.00E+00 | | | |
| Cat 6 | E | 0.00E+00 | 0.00E+00 | | | |
| Dog 1 | E | 0.00E+00 | 1.59E-03 | 0.00E+00 | 3.37E-01 | -1.00E+00 |
| Dog 2 | E | 0.00E+00 | 0.00E+00 | | | |
| Dog 3 | E | 0.00E+00 | 0.00E+00 | | | |
| Ferret 1 | E | 0.00E+00 | 0.00E+00 | | | |
| Hamster 1 | E | 0.00E+00 | 0.00E+00 | | | |
| Hamster 2 | E | 0.00E+00 | 0.00E+00 | | | |
| Hamster 3 | E | 0.00E+00 | 0.00E+00 | | | |
| Cat 1 | M | 0.00E+00 | 0.00E+00 | | | |
| Cat 2 | M | 9.98E-04 | 4.29E-04 | 2.33E+00 | | |
| Cat 3 | M | 9.90E-04 | 0.00E+00 | | | |
| Cat 4 | M | 9.82E-04 | 0.00E+00 | | | |
| Cat 5 | M | 0.00E+00 | 0.00E+00 | | | |
| Cat 6 | M | 0.00E+00 | 0.00E+00 | | | |
| Dog 1 | M | 3.49E-04 | 0.00E+00 | | 4.10E-04 | 4.83E+00 |
| Dog 2 | M | 8.79E-04 | 0.00E+00 | | | |
| Dog 3 | M | 1.89E-04 | 0.00E+00 | | | |
| Ferret 1 | M | 3.88E-04 | 0.00E+00 | | | |
| Hamster 1 | M | 9.35E-04 | 0.00E+00 | | | |
| Hamster 2 | M | 7.85E-04 | 0.00E+00 | | | |
| Hamster 3 | M | 9.05E-04 | 0.00E+00 | | | |
| Cat 1 | N | 0.00E+00 | 1.51E-03 | 0.00E+00 | | |
| Cat 2 | N | 5.16E-04 | 2.10E-03 | 2.46E-01 | | |
| Cat 3 | N | 5.08E-04 | 3.72E-04 | 1.36E+00 | | |
| Cat 4 | N | 5.05E-04 | 8.23E-04 | 6.14E-01 | | |
| Cat 5 | N | 0.00E+00 | 0.00E+00 | | | |
| Cat 6 | N | 0.00E+00 | 0.00E+00 | | 9.54E-02 | -1.81E+00 |
| Dog 1 | N | 3.17E-04 | 1.60E-03 | 1.98E-01 | | |
| Dog 2 | N | 5.11E-04 | 6.76E-04 | 7.55E-01 | | |
| Dog 3 | N | 3.79E-04 | 1.63E-03 | 2.32E-01 | | |
| Ferret 1 | N | 5.16E-05 | 0.00E+00 | | | |
| Hamster 1 | N | 4.90E-04 | 0.00E+00 | | | |

| | | | | | | |
|---|---|---|---|---|---|---|
| Hamster 2 | N | 3.91E-04 | 0.00E+00 | | | |
| Hamster 3 | N | 4.62E-04 | 3.04E-04 | 1.52E+00 | | |
| Cat 1 | orf10 | 0.00E+00 | 0.00E+00 | | | |
| Cat 2 | orf10 | 0.00E+00 | 0.00E+00 | | | |
| Cat 3 | orf10 | 0.00E+00 | 0.00E+00 | | | |
| Cat 4 | orf10 | 0.00E+00 | 0.00E+00 | | | |
| Cat 5 | orf10 | 0.00E+00 | 0.00E+00 | | | |
| Cat 6 | orf10 | 0.00E+00 | 0.00E+00 | | | |
| Dog 1 | orf10 | 0.00E+00 | 0.00E+00 | | | |
| Dog 2 | orf10 | 0.00E+00 | 0.00E+00 | | | |
| Dog 3 | orf10 | 0.00E+00 | 0.00E+00 | | | |
| Ferret 1 | orf10 | 0.00E+00 | 0.00E+00 | | | |
| Hamster 1 | orf10 | 0.00E+00 | 0.00E+00 | | | |
| Hamster 2 | orf10 | 0.00E+00 | 0.00E+00 | | | |
| Hamster 3 | orf10 | 0.00E+00 | 0.00E+00 | | | |
| Cat 1 | orf1ab | 0.00E+00 | 0.00E+00 | | | |
| Cat 2 | orf1ab | 1.04E-04 | 6.47E-05 | 1.60E+00 | | |
| Cat 3 | orf1ab | 9.98E-05 | 9.47E-05 | 1.05E+00 | | |
| Cat 4 | orf1ab | 1.25E-04 | 2.34E-05 | 5.36E+00 | | |
| Cat 5 | orf1ab | 3.92E-05 | 1.78E-05 | 2.20E+00 | | |
| Cat 6 | orf1ab | 2.60E-05 | 0.00E+00 | | | |
| Dog 1 | orf1ab | 1.97E-04 | 3.71E-05 | 5.31E+00 | 1.31E-02 | 2.91E+00 |
| Dog 2 | orf1ab | 1.68E-04 | 1.75E-04 | 9.62E-01 | | |
| Dog 3 | orf1ab | 2.34E-04 | 8.23E-05 | 2.84E+00 | | |
| Ferret 1 | orf1ab | 3.53E-05 | 8.41E-05 | 4.20E-01 | | |
| Hamster 1 | orf1ab | 7.61E-05 | 1.67E-05 | 4.56E+00 | | |
| Hamster 2 | orf1ab | 6.01E-05 | 0.00E+00 | | | |
| Hamster 3 | orf1ab | 1.03E-04 | 3.15E-05 | 3.27E+00 | | |
| Cat 1 | orf3a | 0.00E+00 | 0.00E+00 | | | |
| Cat 2 | orf3a | 0.00E+00 | 0.00E+00 | | | |
| Cat 3 | orf3a | 0.00E+00 | 0.00E+00 | | | |
| Cat 4 | orf3a | 0.00E+00 | 0.00E+00 | | | |
| Cat 5 | orf3a | 0.00E+00 | 0.00E+00 | | | |
| Cat 6 | orf3a | 0.00E+00 | 0.00E+00 | | | |
| Dog 1 | orf3a | 0.00E+00 | 0.00E+00 | | 3.37E-01 | 1.00E+00 |
| Dog 2 | orf3a | 5.94E-04 | 0.00E+00 | | | |
| Dog 3 | orf3a | 0.00E+00 | 0.00E+00 | | | |
| Ferret 1 | orf3a | 0.00E+00 | 0.00E+00 | | | |
| Hamster 1 | orf3a | 0.00E+00 | 0.00E+00 | | | |
| Hamster 2 | orf3a | 0.00E+00 | 0.00E+00 | | | |
| Hamster 3 | orf3a | 0.00E+00 | 0.00E+00 | | | |
| Cat 1 | orf6 | 0.00E+00 | 0.00E+00 | | 9.82E-02 | 1.79E+00 |

| | | | | | | |
|---|---|---|---|---|---|---|
| Cat 2 | orf6 | 0.00E+00 | 0.00E+00 | | | |
| Cat 3 | orf6 | 4.11E-04 | 0.00E+00 | | | |
| Cat 4 | orf6 | 0.00E+00 | 0.00E+00 | | | |
| Cat 5 | orf6 | 0.00E+00 | 0.00E+00 | | | |
| Cat 6 | orf6 | 0.00E+00 | 0.00E+00 | | | |
| Dog 1 | orf6 | 0.00E+00 | 0.00E+00 | | | |
| Dog 2 | orf6 | 0.00E+00 | 0.00E+00 | | | |
| Dog 3 | orf6 | 0.00E+00 | 0.00E+00 | | | |
| Ferret 1 | orf6 | 7.88E-04 | 0.00E+00 | | | |
| Hamster 1 | orf6 | 0.00E+00 | 0.00E+00 | | | |
| Hamster 2 | orf6 | 0.00E+00 | 0.00E+00 | | | |
| Hamster 3 | orf6 | 9.13E-04 | 0.00E+00 | | | |
| Cat 1 | orf7a | 0.00E+00 | 0.00E+00 | | | |
| Cat 2 | orf7a | 0.00E+00 | 0.00E+00 | | | |
| Cat 3 | orf7a | 0.00E+00 | 0.00E+00 | | | |
| Cat 4 | orf7a | 0.00E+00 | 0.00E+00 | | | |
| Cat 5 | orf7a | 0.00E+00 | 0.00E+00 | | | |
| Cat 6 | orf7a | 0.00E+00 | 0.00E+00 | | | |
| Dog 1 | orf7a | 0.00E+00 | 0.00E+00 | | 3.37E-01 | 1.00E+00 |
| Dog 2 | orf7a | 0.00E+00 | 0.00E+00 | | | |
| Dog 3 | orf7a | 4.68E-04 | 0.00E+00 | | | |
| Ferret 1 | orf7a | 0.00E+00 | 0.00E+00 | | | |
| Hamster 1 | orf7a | 0.00E+00 | 0.00E+00 | | | |
| Hamster 2 | orf7a | 0.00E+00 | 0.00E+00 | | | |
| Hamster 3 | orf7a | 0.00E+00 | 0.00E+00 | | | |
| Cat 1 | orf8 | 0.00E+00 | 0.00E+00 | | | |
| Cat 2 | orf8 | 0.00E+00 | 0.00E+00 | | | |
| Cat 3 | orf8 | 4.81E-04 | 0.00E+00 | | | |
| Cat 4 | orf8 | 0.00E+00 | 0.00E+00 | | | |
| Cat 5 | orf8 | 0.00E+00 | 0.00E+00 | | | |
| Cat 6 | orf8 | 0.00E+00 | 0.00E+00 | | | |
| Dog 1 | orf8 | 1.78E-03 | 1.81E-03 | 9.83E-01 | 9.18E-01 | 1.05E-01 |
| Dog 2 | orf8 | 6.28E-04 | 2.77E-03 | 2.27E-01 | | |
| Dog 3 | orf8 | 2.02E-03 | 0.00E+00 | | | |
| Ferret 1 | orf8 | 0.00E+00 | 0.00E+00 | | | |
| Hamster 1 | orf8 | 0.00E+00 | 0.00E+00 | | | |
| Hamster 2 | orf8 | 0.00E+00 | 0.00E+00 | | | |
| Hamster 3 | orf8 | 0.00E+00 | 0.00E+00 | | | |
| Cat 1 | S | 9.08E-06 | 0.00E+00 | | | |
| Cat 2 | S | 6.16E-04 | 0.00E+00 | | | |
| Cat 3 | S | 6.12E-04 | 0.00E+00 | | 2.27E-04 | 5.19E+00 |
| Cat 4 | S | 4.49E-04 | 0.00E+00 | | | |
| Cat 5 | S | 3.40E-05 | 0.00E+00 | | | |

| | | | | |
|---|---|---|---|---|
| Cat 6 | S | 4.76E-05 | 0.00E+00 | |
| Dog 1 | S | 2.33E-04 | 0.00E+00 | |
| Dog 2 | S | 6.08E-04 | 0.00E+00 | |
| Dog 3 | S | 2.96E-04 | 0.00E+00 | |
| Ferret 1 | S | 1.77E-04 | 1.07E-04 | 1.66E+00 |
| Hamster 1 | S | 5.00E-04 | 0.00E+00 | |
| Hamster 2 | S | 5.25E-04 | 0.00E+00 | |
| Hamster 3 | S | 4.66E-04 | 0.00E+00 | |
